# Transition to chaos separates learning regimes and relates to measure of consciousness in recurrent neural networks

**DOI:** 10.1101/2024.05.15.594236

**Authors:** Dana Mastrovito, Yuhan Helena Liu, Lukasz Kusmierz, Eric Shea-Brown, Christof Koch, Stefan Mihalas

**Affiliations:** Allen Institute; University of Washington; Allen Institute for Brain Science

## Abstract

Recurrent neural networks exhibit chaotic dynamics when the variance in their connection strengths exceed a critical value. Recent work indicates connection variance also modulates learning strategies; networks learn ”rich” representations when initialized with low coupling and ”lazier”solutions with larger variance. Using Watts-Strogatz networks of varying sparsity, structure, and hidden weight variance, we find that the critical coupling strength dividing chaotic from ordered dynamics also differentiates rich and lazy learning strategies. Training moves both stable and chaotic networks closer to the edge of chaos, with networks learning richer representations before the transition to chaos. In contrast, biologically realistic connectivity structures foster stability over a wide range of variances. The transition to chaos is also reflected in a measure that clinically discriminates levels of consciousness, the perturbational complexity index (PCIst). Networks with high values of PCIst exhibit stable dynamics and rich learning, suggesting a consciousness prior may promote rich learning. The results suggest a clear relationship between critical dynamics, learning regimes and complexity-based measures of consciousness.

---

As learning in artificial networks continues to amass practical successes, theorists have been making significant strides in rigorously characterizing the behavior of these models and explaining why they perform so well [1–8]. These theoretical tools present an opportunity to unravel mysteries in biological neural networks [9], such as how the learning rule and/or initial priors of a network could alter its learning dynamics and representations [10–13]. Among various theoretical developments that contribute to this progress, a popular theme is that networks can be successfully trained to learn a task using two distinct strategies: rich learning and lazy learning [14–23]. Interpolation between the two learning regimes can be achieved through adjusting the variance in connection strengths at initialization [14, 15, 24]. This adjustment tunes the extent to which the network alters its internal representation to fit the task statistics, leading to various degrees of task-specific representations. Consequently, whether learning occurs in the rich or lazy regime can have a profound effect on the nature of what the network learns and its performance in unseen situations post-training [15, 16].

Connection strength in artificial networks has also been studied in the context of dynamical systems theory, where it has been shown that networks exhibit a transition between ordered and chaotic dynamics when the variance of their connection strengths exceed a critical value [25]. The transition point between ordered and chaotic regimes can be identified mathematically using the maximum Lyapunov exponent *λ*, which when positive, indicates chaotic dynamics. At intermediate connection strengths, at a transition point in phase space known as the edge of chaos, systems are known to have optimal computational performance, exhibiting maximal information transfer [26, 27], and memory capacity [28]. Although, more recent work has suggested that networks can continue to perform well in the weakly chaotic regime, as degradation in autocorrelation of activity occurs more slowly in the chaotic than non-chaotic regime [29]. Building on these theoretical insights, we vary initial connection strengths of recurrent neural networks (RNNs) and characterize their learning properties, finding a direct correspondence between the transition to chaos, when the largest Lyapunov exponent prior to training becomes positive, and a shift between rich and lazy learning strategies.

The brain has long been theorized to operate at the edge of chaos due in part to the aforementioned optimal properties associated with this dynamical regime. Beyond arguments of optimality, however, it is reasonable to theorize that the highly recurrent brain operates within this regime, as self-organized criticality is observed in many natural systems and recent theory shows that suppression of chaos may be inherent in systems utilizing integrative feedback [30]. In fact, several recent studies have uncovered evidence to suggest that edge of chaos dynamics may underlie the capacity for consciousness itself [31–34]. Although consciousness is difficult to define and measure, one recently developed metric called the perturbational complexity index (PCI) has emerged as a reliable correlate of consciousness where it has been demonstrated to distinguish between brain states (awake, anesthetized, under the influence of psychedelics), and to reflect the potential for recovery in patients with disorders of consciousness [35–37]. The PCI metric is predicated on the theory that the capacity for consciousness relies on the ability to integrate information, and that this ability is achieved through the complex patterns of causal interactions between neurons. However, the original PCI metric is only applicable to systems for which one can employ transcranial magnetic stimulation in combination with electroencephalograpy (EEG). Therefore, in this study, we make use of an estimate of PCI (PCIst) [38] which is more broadly applicable. The metric similarly quantifies the spatiotemporal complexity of the propagation of evoked activity in response to an externally driven perturbation above that of baseline activity. Intuitively, the metric combines spatial principal component analysis (sPCA) with recurrence quantification analysis (RQA) [31, 39], which quantifies the temporal complexity as the recurrences of the evoked dynamics. RQA is commonly used in the analysis of dynamical systems [40] to identify state transitions and has properties that are directly related to Lyapunov exponents [40, 41]. Such methods have successfully been applied to the analysis of biological systems [42–44] where the direct computation of the Lyapunov spectrum is impractical. Nevertheless, no previous study has examined the relationship between PCIst and Lyapunov exponents. We therefore computed PCIst on RNNs and found that it increases as a function of initial connection strength up to the edge of chaos, where it is maximal, and subsequently sharply decreases.

Finally, we compare Lyapunov exponents, learning regime and PCIst on network models initialized with Gaussian weight distributions to those with biologically-realistic connectivity structures at two different scales: that of a cortical column within mouse primary visual cortex and of the mouse whole-brain mesoscopic connectivity. We find that biologically realistic connectivity yields non-chaotic dynamics, where PCIst is high and rich learning is favored, over a wider range of connectivity strengths.

## Summary of results and contributions

Building on three lines of literature — rich versus lazy learning, dynamical systems theory and consciousness — we find and characterize two regimes associated with the initial hidden weight gains below and above the critical point at which networks begin to exhibit chaotic dynamics. Below the critical point, in the ordered learning regime, networks learn rich low-dimensional representations. Beyond the critical point, in the chaotic learning regime, networks gradually transition towards a high-dimensional, lazy learning strategy that is more sensitive to noise. Importantly, we find that models in the chaotic regime, with gains close to the transition, still perform the task with high accuracy, and converge more quickly than in the ordered regime. Further increases in initial connection strength variance results in chaotic dynamics, drastically reducing learning ability. Connection sparsity allows stable dynamics with larger variances, and training moves networks closer to the edge of chaos. Intriguingly, we find that PCIst increases as a function of gain below the transition point, is maximal at the edge of chaos and subsequently sharply decreases. Finally, we show that several key features of biologically-realistic connectivity strongly bias networks towards ordered dynamics and rich learning. We focus on RNNs due to their widespread usage in brain modeling [24, 45–64]. Although in artificial learning networks, our results show a strong relationship between critical dynamics, learning and measures of consciousness.

## 1 Results

In a series of RNNs with systematically varied sparsity and degree of small-world structure, we characterize 1) their dynamical regime (ordered vs chaotic) 2) their PCIst scores and 3) their learning regime (rich vs lazy) as a function of an initial scaling parameter (*g*) on the strength of their hidden weight connectivity (*H*). Specifically, we trained a series of RNNs to perform the ten digit (0-9) sequential MNIST task, in which the network receives a row of pixels from a digit image at each time step. All networks consisted of an input layer, a single recurrent layer with 198 neurons, and a linear readout layer. The hidden layer structure of each network was initialized according to a Watts-Strogatz [65] connectivity rule over a range of nearest neighbors (nNN: 4, 8, 16, 28, 32, 64, 128, 198 = fully connected), and rewiring probabilities (p: 0.0, 0.2, 0.5, 0.8, 1.0), while self-connections were prohibited. Thus, the resulting networks systematically varied in both their degree of sparsity with nNN = 4 most sparse; nNN = 198 least sparse and their degree of small-world structure with p = 0.0 highly structured; p = 1.0 Erdös-Rényi random connectivity (Figure 1). (see Table 1 for number of parameters in each model). Initial hidden layer weights were sampled from a Gaussian distribution with mean *µ* = 0 and standard deviation 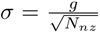 where gain (g : 0.5, 0.75, 1.0, 1.25, 1.5, 1.75, 2.0, 2.25, 2.5, 2.75, 3.0, 5.0) and *N_nz_* is the number of non-zero elements, such that variance is scaled in accordance with sparsity. Network sparsity was maintained over training by restricting training to only the non-zero elements of the hidden weight matrices.

**Figure 1:**
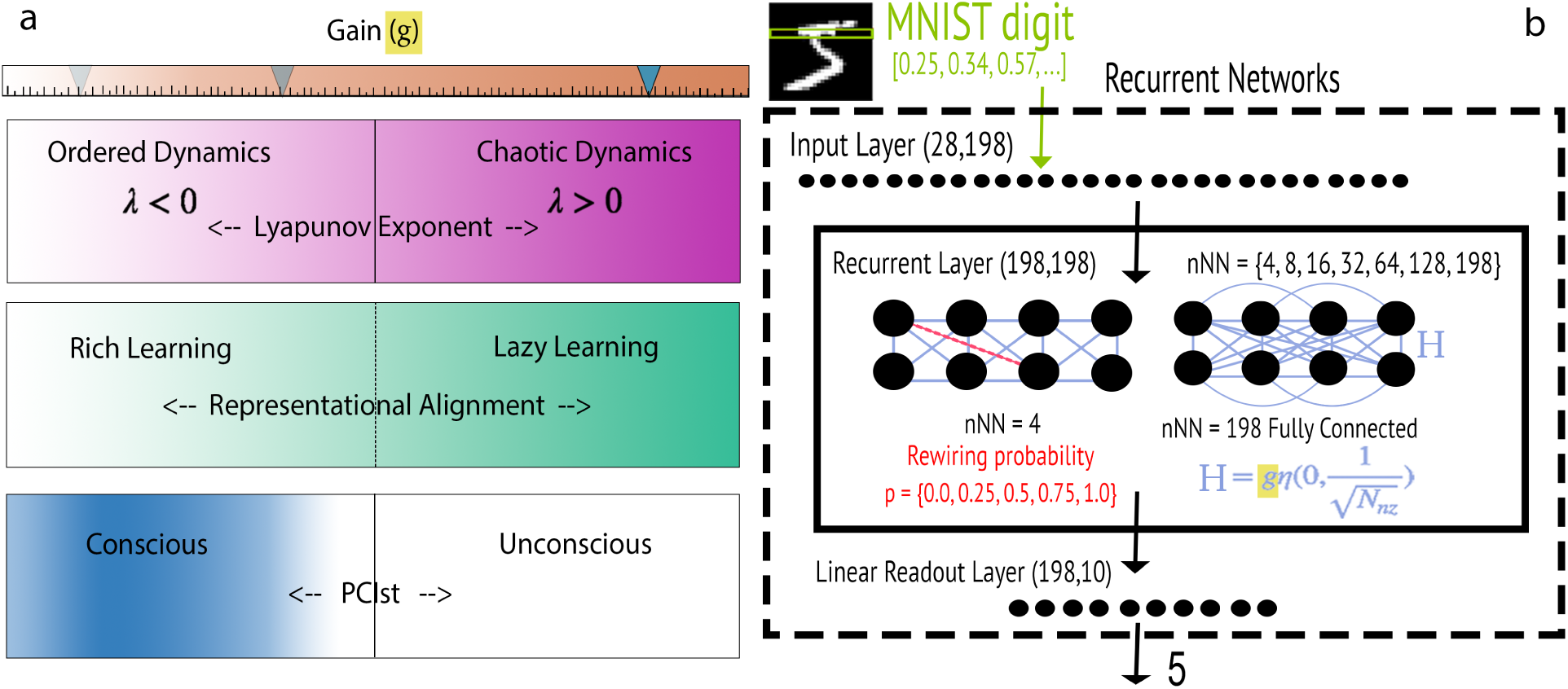
Summary of Results. **(a)** A single gain parameter (g) scaling initial hidden weights modulates ordered and chaotic dynamical regimes, rich and lazy learning strategies as well as a measure associated with consciousness. **(b)** Model Construction. Models consist of an input layer of size 28, a single recurrent layer of 198 hidden units. The hidden layer connectivity matrix (*H*) is initialized as a Watts Strogatz network with number of nearest neighbor connections nNN = *{*4, 8, 16, 28, 32, 64, 128, 198 = fully connected*}* and rewiring probability p = *{*0.0, 0.2, 0.5, 0.8, 1 = Erdös-Rényi*}*. Initial hidden layer connection strengths are sampled from a normal distribution 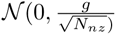, where *N_nz_* is the number of nonzero elements and gain g = 0.5, 0.75, 1.0, 1.25, 1.5, 1.75, 2.0, 2.25, 2.5, 2.75, 3.0, 5.0. The location of zero/non-zero elements are maintained over training on the sMNIST task.

**Table 1:**
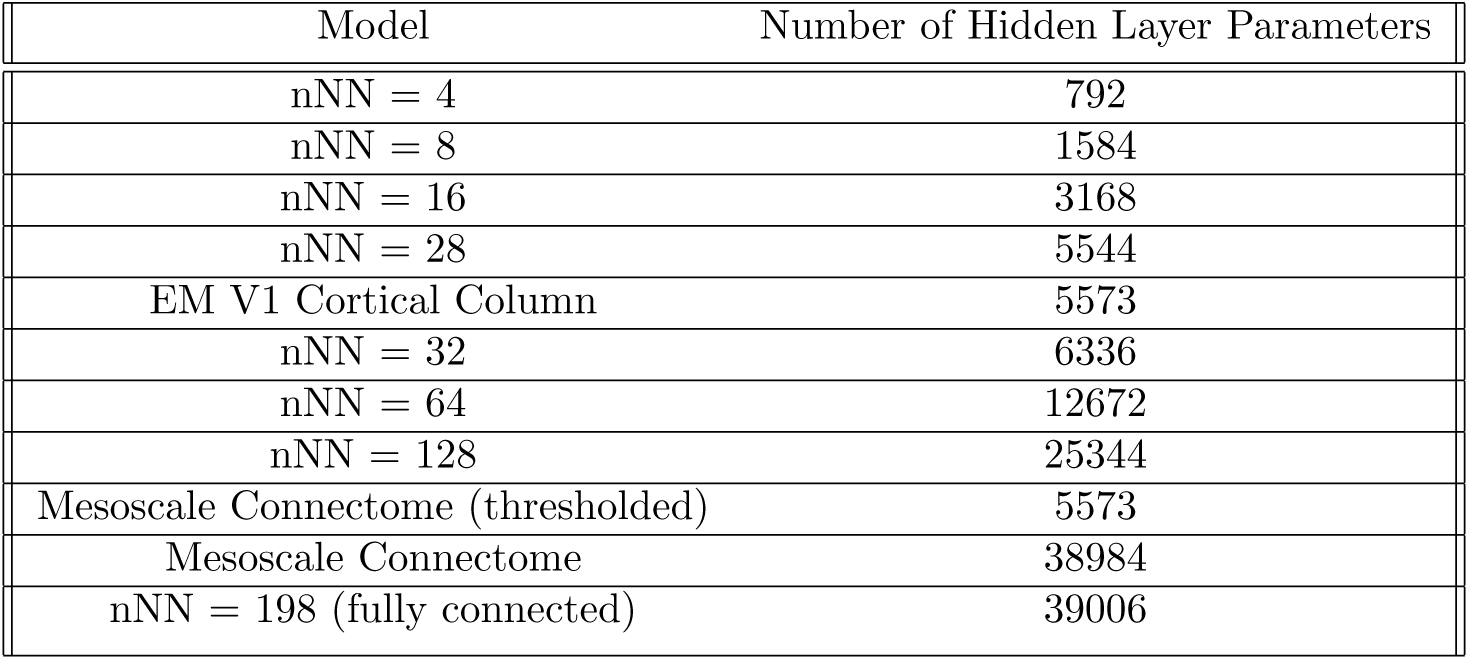
Model Hidden Weight Sparsity and Parameter Count.

### 1.1 Chaotic and Ordered Learning

We computed finite time Lyapunov spectra for each RNN model as the eigenvalues of the product of the Jacobians along the data-driven network trajectory, averaged over a batch of sMNIST test data [66] (see section 2.3.1 Lyapunov Exponents in Methods for details). Chaotic dynamics are indicated when the largest Lyapunov exponent *λ >* 0. For models at each sparsity level (nNN), we define the transition point between ordered and chaotic learning regimes in terms of a single multiplicative parameter *g*, the gain in connection strength of the initial (pre-training) hidden layer connectivity. Specifically, we define the critical transition point associated with each model’s degree of sparsity as the gain (*g_cnNN_*) at which the initialized (prior to training) model’s largest Lyapunov exponent *λ* becomes positive. We find that the transition from ordered to chaotic learning regimes shifts towards lower values of gain (*g*) as models become less sparse (nNN increases)(Figure 2a,b). Rewiring probability, which shifts network structure away from modular small-world structures towards Erdös-Rényi random connectivity as it increases, had a much smaller impact on network dynamics than did the sparsity. In fact, we found that the transition point often did not vary substantially with rewiring probability (Table S1, Figure 2a). Therefore, unless otherwise noted, throughout the manuscript we generally report results for rewiring probability of 1.0 = Erdös-Rényi, as it is the most commonly used network initialization strategy, but see Figure S1 for similar results obtained with rewiring probabilities p: 0.0, 0.2, 0.5, 0.8. Consistent with prior work [27, 67, 68], we found that networks self-tuned towards criticality as a result of training with back-propagation, such that the maximum Lyapunov exponents after training shifts closer to zero for all models (Figure 3a). However, we find that changes in the magnitude of the maximum Lyapunov exponent over training decreases substantially as the models enter the chaotic regime (Figure 3b), suggesting that models are less able to tune towards the edge of chaos if the variance in their initial connectivity strength is too large.

**Figure 2:**
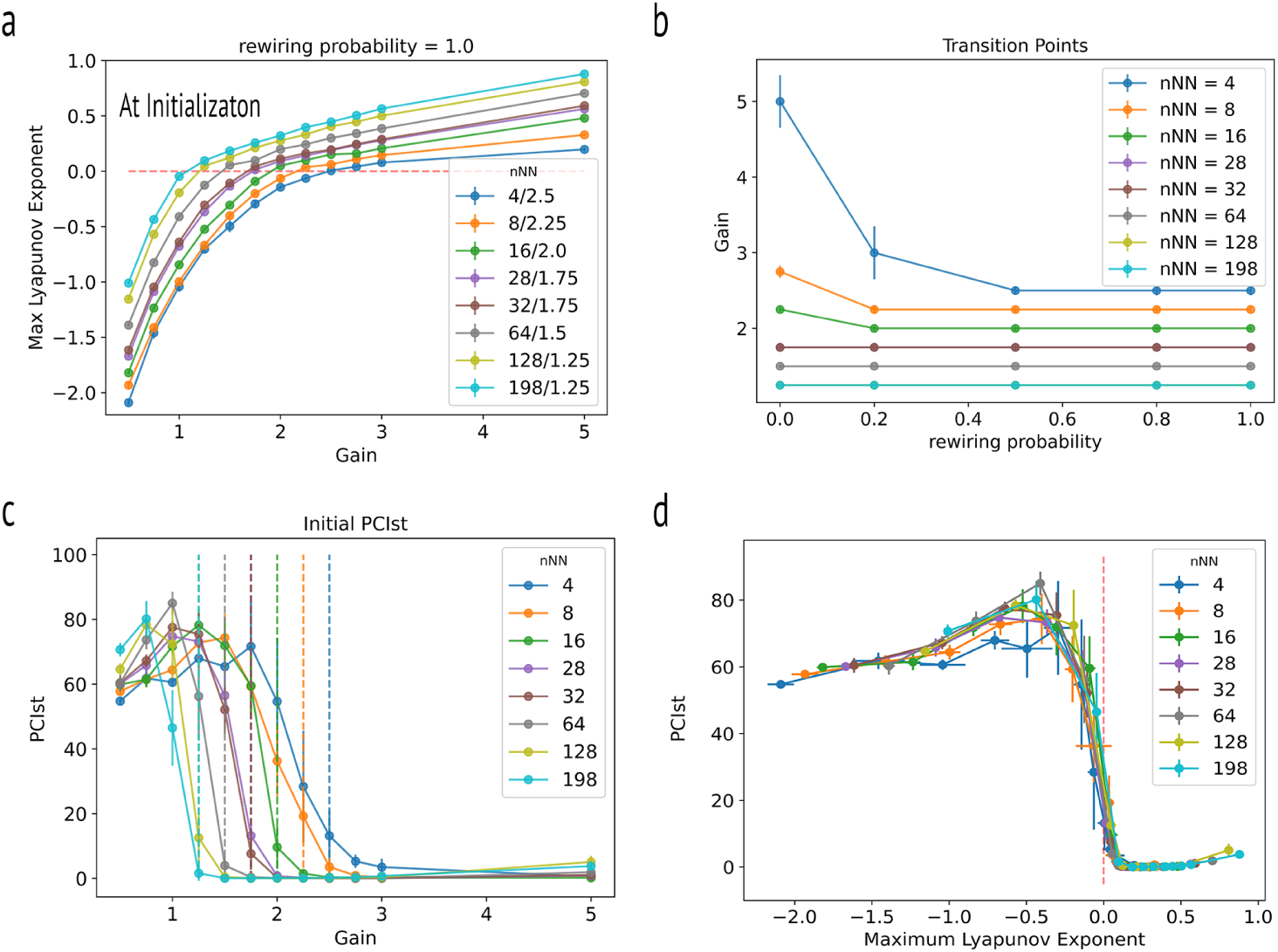
PCIst and Maximum Lyapunov Exponents Before Training **(a)** Maximum Lyapunov Exponent *λ* for driven models with rewiring probability *p* = 1.0 along trajectory. Gain at which each network transitions from ordered to chaotic regime *g_cnNN_* is the gain at which *λ* becomes positive for each level of sparsity nNN. **(b)** Model Transition points. Transitions to chaos shift to smaller gains as hidden layer sparsity increases. **(c)** PCIst in pre-trained networks increases as a function of gain and begins to decrease just before *g_cnNN_* where it decreases sharply. **(d)** PCIst in pre-trained networks begins to decrease as the maximum Lyapunov exponent *λ* approaches zero.

**Figure 3:**
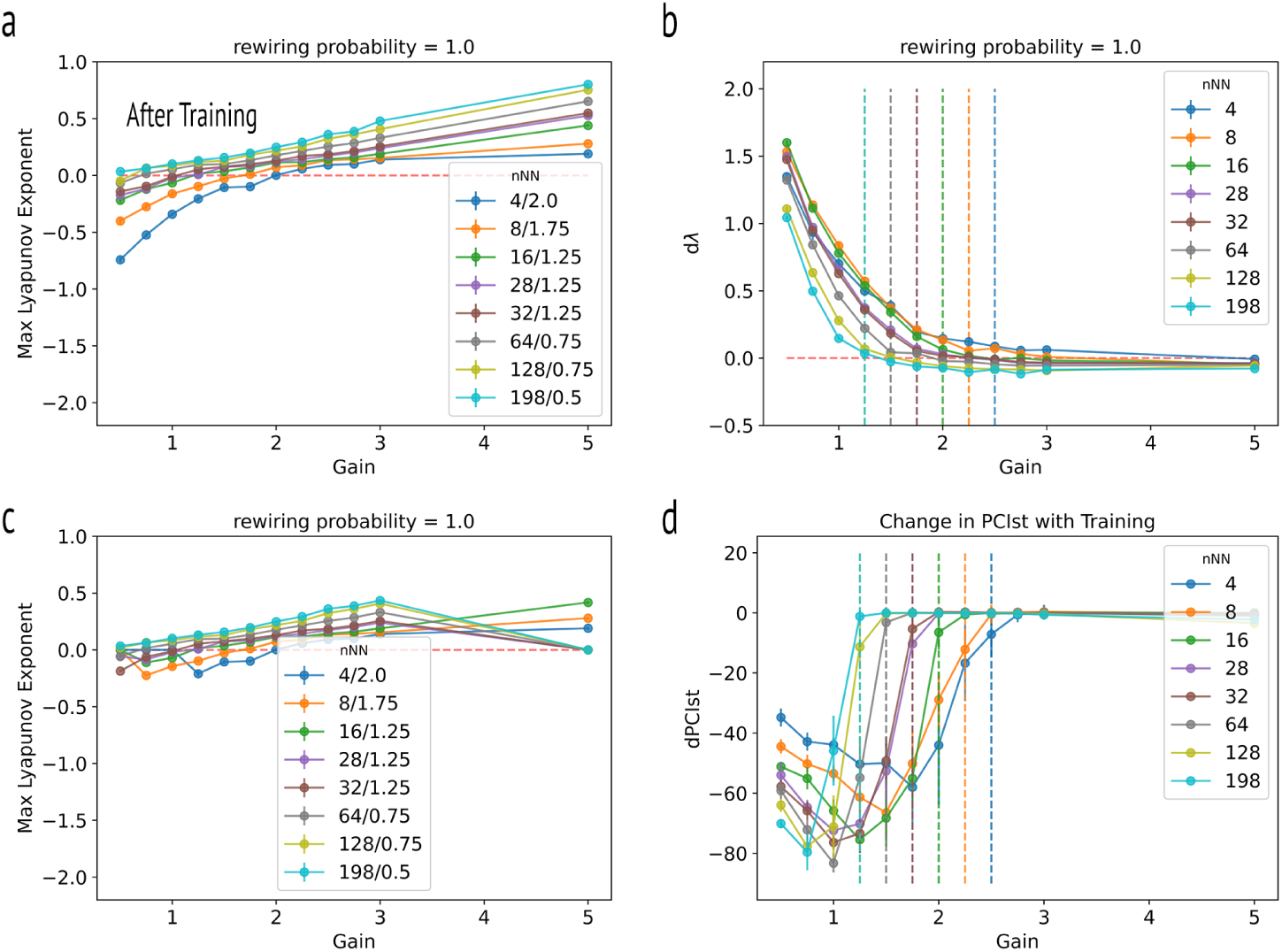
PCIst and Maximum Lyapunov Exponents After Training **(a)** Maximum Lyapunov Exponent *λ* after training for 100 epochs. Models tune towards the edge of chaos with training. **(b)** Change in maximum Lyapunov exponent *λ* with training (100 epochs). Models trained in a highly chaotic regime exhibit smaller changes in *λ* with training. **(c)** Maximum Lyapunov exponent in fully-trained models, trained till 90% accuracy. **(d)** Changes in PCIst over training decrease in the chaotic regime.

Although models in the weakly chaotic regime are often able to obtain good accuracy (See section 1.2 for model performance), the trained model dynamics in the ordered and chaotic regimes are qualitatively very different. To illustrate the difference in model solution space, we project the hidden state space of the trained model responses to sMNIST test samples into 2-dimensions (for ease of visualization) using principle component analysis (PCA), and find qualitatively different solutions to the task are learned on either side of *g_cnNN_*. In the ordered learning regime, models exhibit a smooth trajectory from initial state (at the origin) towards a final state located in a cluster associated with digit identity. In contrast, in the chaotic learning regime, just past the transition, where models nevertheless still learn to perform the task, models exhibit a jagged trajectory (Figure S2, illustrating the resulting chaotic dynamics).

Finally, we tested for model sensitivity to noise at test time. Models, trained without noise were subjected to additional Gaussian noise during testing (See Methods for details). Because small perturbations should be amplified in the chaotic regime, we expected that models in the chaotic learning regime would be more sensitive to injected noise. Indeed we found that accuracy dropped more substantially for models in the chaotic than ordered regimes when exposed to additional Gaussian noise *N* (0,.01) at test time (Figure 4c).

**Figure 4:**
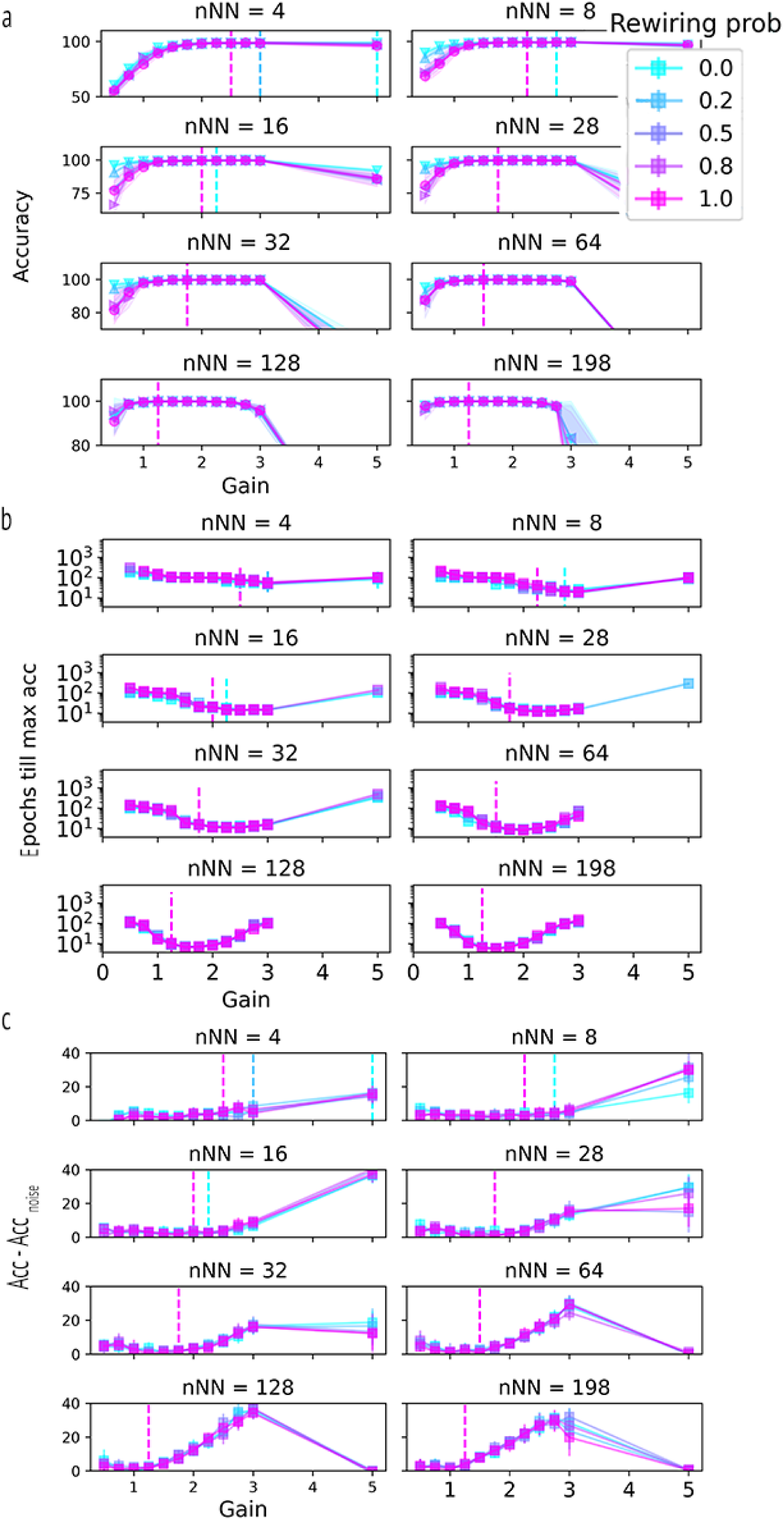
Model Performance. **(a)** Accuracy after 100 epochs. Dashed lines indicates *g_cnNN_*, where models achieve their highest accuracy. As sparsity decreases ability to learn the task falls sharply as gain increases beyond ≥ 3.0. **(b)** Number of Epochs till 90% accuracy. **(c)** Difference in accuracy with additional noise during testing. Networks are more sensitive to noise in the chaotic regime.

### 1.2 Model Performance

Model performance was assessed via both the test accuracy achieved as well as speed of learning. Test accuracy was quantified as the average percent correct over all sMNIST test samples after 100 epochs of training, while speed of learning was assessed by the number of epochs required to reach at least 90% accuracy. For both metrics, the reported values are the mean over 10 model instantiations with the same model parameters: nNN, rewiring probability (p), gain (g). The least sparse models achieved higher accuracy after 100 epochs in comparison to sparse ones; while performance on models with equal sparsity, differing only in connectivity structure, performed nearly identically. After 100 epochs of training, accuracy increased as a function of gain up to *g_cnNN_*, where all models achieved their highest accuracy (Figure 4a). Of critical interest is the fact that models continued to have high accuracy after the transition into the chaotic learning regime, while also learning faster in this regime (Figure 4b), with small increases in initial weight gain above *g_cnNN_*. We are not the first to report that models can perform well in the weakly chaotic regime. This result is consistent with previously reported numerical experiments on the lazy training regime [14, 69] where it was also found that lazy models converged faster. As we will show in the subsequent sections, however, learning in this regime occurs using a ”lazier” strategy consistent with previous work indicating that dimensionality expansion in the weakly chaotic regime allows for linearly separable representations at the readout layer [70]. This is can be seen in Figure 5a,e, where the dimensionality of the trained networks at higher gains increases and is reflected by greater weight changes in the readout layer in successfully trained chaotic networks. However, further increases in gain result in sharp decreases in accuracy (Figure 4a). Again, analogously to the dynamical regime, we found that hidden layer sparsity had a much greater effect on model performance than connectivity structure varied through rewiring probability.

**Figure 5:**
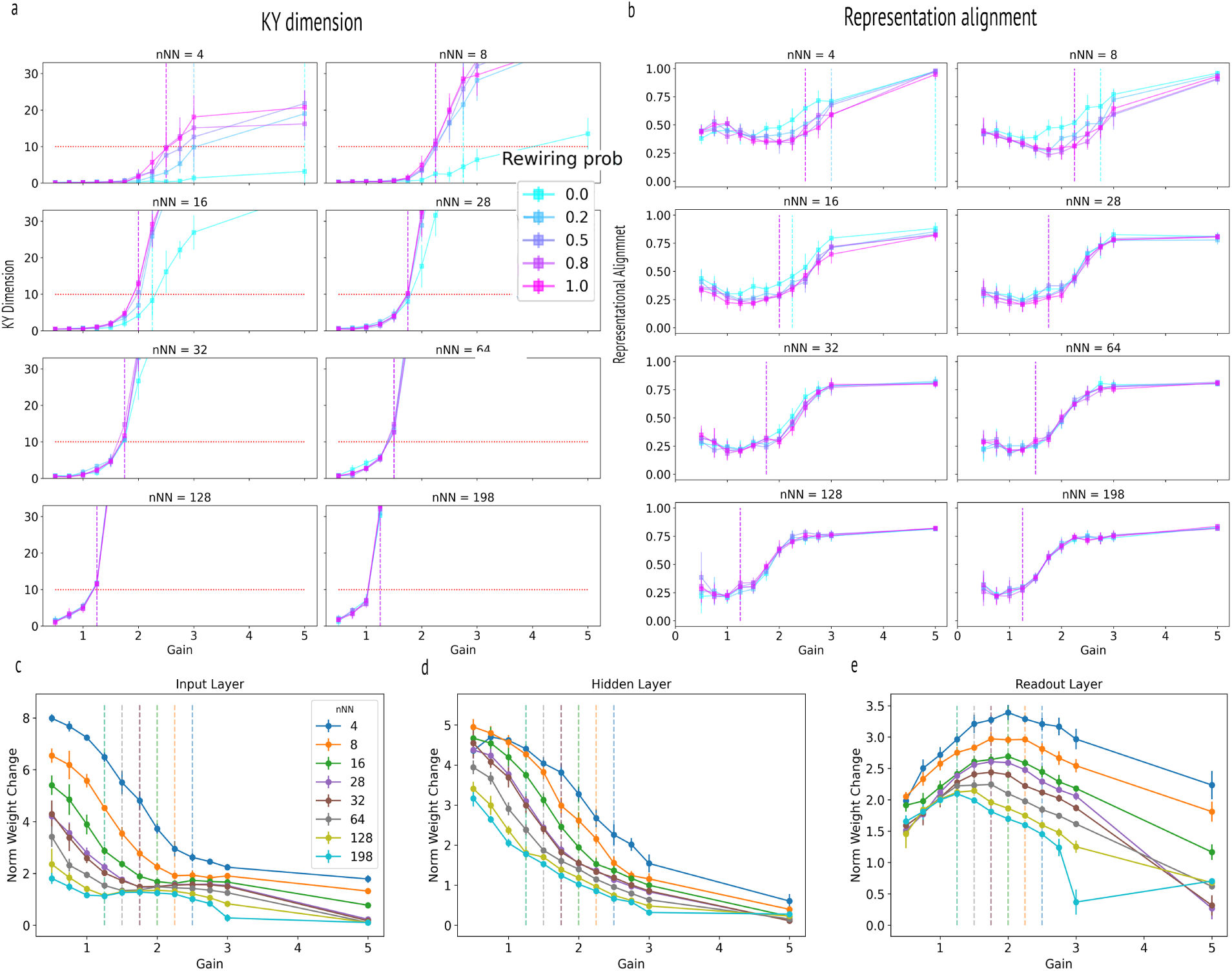
Rich vs. Lazy Learning **(a)** Kaplan-Yorke (KY) dimensionality. Dashed lines indicate *g_cnNN_*. **(b)** Representation alignment for different sparsity and rewiring probabilities. **(c)** Norm Weight Change of input layer weights over training. **(d)** Norm weight change of hidden layer. **(e)** Norm weight change of readout layer. Weight changes in the input and hidden layers decrease smoothly with gain. In the readout layer, however, weight changes are non-monotonic, increasing as a function of gain up to *g_cnNN_* and then decreasing.

### 1.3 Rich vs Lazy Learning Regime

In the following sections, we report several metrics used to characterize the learning regime. First, we computed the Kaplan-Yorke (KY) dimension of the trained models derived from the Lyapunov spectrum. It has previously been demonstrated that, with random initialization, lazy learning leads to higher dimensional task-agnostic representation, whereas rich learning results in lower dimensional task-specific representations [12, 69]. Second, lazier learning results in less modification to the hidden weight parameters [14, 24, 69], and we report the vector norm of the magnitude weight changes at the input, hidden, and output layers. Finally, we compute representation alignment, which quantifies the directional change in a representational similarity matrix before and after training; lazier learning should result in higher representation alignment [16]. See also Supplementary Results where we also quantify the directional shift in the neural tangent kernel (NTK) pre- and post-training (Figure S4). This measure, referred to as tangent kernel alignment, provides another method for quantifying the richness/laziness of learning [16]. We emphasize that in this study, laziness and richness are quantified on a continuum rather than categorically, with lazier learning corresponding to smaller network changes to achieve learning of a task. In other words, we adopt the notion of an effective learning regime used in [71], which gauges effective richness or laziness by post-training changes rather than on initialization.

#### 1.3.1 Dimensionality

We computed the Kaplan-Yorke (KY) dimension of the trained models derived from the Lyapunov spectrum (see section 2.6 in Methods for details). The KY dimensionality increased as a function of gain, accompanied by a shift towards lazier learning. Notably, models learned very low-dimensional solutions at the smallest gains (Figure 5a). Interestingly, as the task consists of 10 classes, the dimensionality of the trained models exceeded 10 at *g_cnNN_*, for all but the sparsest and most small-world models. KY dimensionality continued to increase with further increases in gain. We trained an additional series of models for rewiring probability = 1.0 only, on the 2-digit sMNIST task, using digits [2,5] and found that models exceeded KY dimension of 2 rather than 10 at *g_cnNN_* (Figure S3).

#### 1.3.2 Weight Changes

We found that the norm of weight changes over training in both the input and hidden layers decrease smoothly and monotonically as a function of gain for all models (Figure 5c-d). However, in the readout layer, changes in weights increase as models approach *g_cnNN_*, after which it gradually begins to decrease. This is consistent with previous work that found lazy learning can result from expanding the dimensionality of input signals. Specifically, random projections to the hidden layer create a representation that facilitates linear separability. Consequently, learning primarily occurs in the readout weights [72]. So as gain increases, learning smoothly shifts away from rich hidden layer representations, as evidenced by large magnitude changes in the hidden layer and when models engage in lazy learning, learning is confined to the readout layer. As the gain increases far into the chaotic regime where models fail to learn the task, presumably due to numeric instability, the normed weight changes of all layers approach zero.

#### 1.3.3 Representation Alignment

As expected, we find that representation alignment (see Methods section 2.7.2) between trained and untrained models largely increases with increased gain, indicating greater laziness as gain increases (Figure 5b). Although the curves are not always monotonic, the representation alignment typically increases just before the *g_cnNN_* transition.

### 1.4 PCIst

PCIst was assessed in all networks before and after training on the sMNIST task. PCIst at initialization increased as a function of gain across all network architectures with g *≤* 1, however initial PCIst begins to decrease as the gain approaches *g_cnNN_* for all models and nears zero when models enter the chaotic regime (Figure 2c). In section 1.1 we found that the gain at which the maximum Lyapunov exponent becomes positive shifts to ever smaller gains as sparsity decreases. Consistent with the maximum Lyapunov exponent itself, the gain at which PCIst begins to decrease also shifts to smaller gains as sparsity decreases. The metric hits a maximum prior to the critical point, as models near the edge of chaos transition (Figure 2d). Critically, PCIst reaches its minimum at *g_cnNN_* or, in the sparser models, just beyond the point at which models become chaotic. Also consistent with the maximum Lyapunov exponent, PCIst decreases with learning in models initialized in a non-chaotic regime, as the models tune towards towards criticality as a result of training with back-propagation. For both the maximum Lyapunov exponent and PCIst, we do not observe decreases as a result of training when initialized in the chaotic regime (Figure 3b,d). All models considered here were of equal size. To test for finite size effects, we created two larger models with 500 and 1000 hidden units (Figure S12) and found that the point at which PCIst begins to decrease moves closer to the edge of chaos as networks increase in size. In all cases, we find PCIst is non-zero in the ordered regime where richer learning strategies are favored and maximal just before the transition to chaos.

### 1.5 Biologically Realistic Connectivity

Biological brain networks are known to have small-world architectures [65, 73], but differ from the models previously explored in important ways including the distribution of their weights (Figure 7a, the degree distribution of each neuron, and adherence to Dale’s law. We therefore trained a series of models with hidden weight matrices defined by biologically realistic connectivity structures at two different scales. The first connectivity structure was defined by the normalized whole-brain mesoscopic connectivity density of the mouse connectome [74, 75] as measured from hundreds of anterograde tracing experiments. Normalized connection density is defined as the directional inter-regional connection strength divided by the product of the size of the regions. The mesoscopic connectivity model has a similar number of parameters as the nNN = 198 (fully connected) model, but the distribution of connection strengths has a longer-tailed distribution (Figure 7a). As connection strengths are by definition all positive, for these models, an equal number of positive and negative weights were assigned randomly. The second model, at the scale of a single microcolumn, was derived from electron microscopy of the mouse primary visual cortex (V1)[]. In this model, the connection strength *H_ij_* from neuron j to neuron i was defined as the sum of the volume of synaptic densities at neuron i coming from neuron j. For comparison, this cortical column model has approximately the same number of parameters as the nNN = 28 models (See Table 1). The sign of each connection weight is cell-type specific as dictated by Dale’s Law, such that all post-synaptic connections of excitatory neurons are positive, and visa-versa. We now describe the differences we observed in model performance, dynamical and learning regimes in biologically realistic connectivity structures in contrast to models initialized with Gaussian hidden weight distributions previously described.

#### 1.5.1 Chaotic and Ordered Dynamics

In the case of the mesoscopic connectivity model, we find that the relationship between initial hidden weight gain and the maximum Lyapunov exponent pre-training has a shallower slope, such that it enters the chaotic regime at a higher gain than the fully connected Gaussian model (Figure 6a:green dash-dot/cyan). Additionally, the magnitude of the largest Lyapunov exponent remains below that of the fully connected model at higher gain values. For comparison to the cortical column model, we also trained a modified mesocopic model with a thresholded connectivity structure, such that sparsity was matched to the cortical column model. Here again, relative to the nNN = 28 Gaussian model, we see that the mesoscopic model has a shallower slope and the magnitude of the largest Lyapunov exponent remains below that of the Gaussian model at high gains (Figure 6a:dark blue dashed/brown), suggesting that the long-tailed distribution of the weights of the mesoscopic connectivity structure affords the model greater stability than a Gaussian model of equal sparsity. See also Figures S6, S7 for comparison of eigenspectrum and Lypaunov spectrums for Gaussian and biologically realistic connectivity.

**Figure 6:**
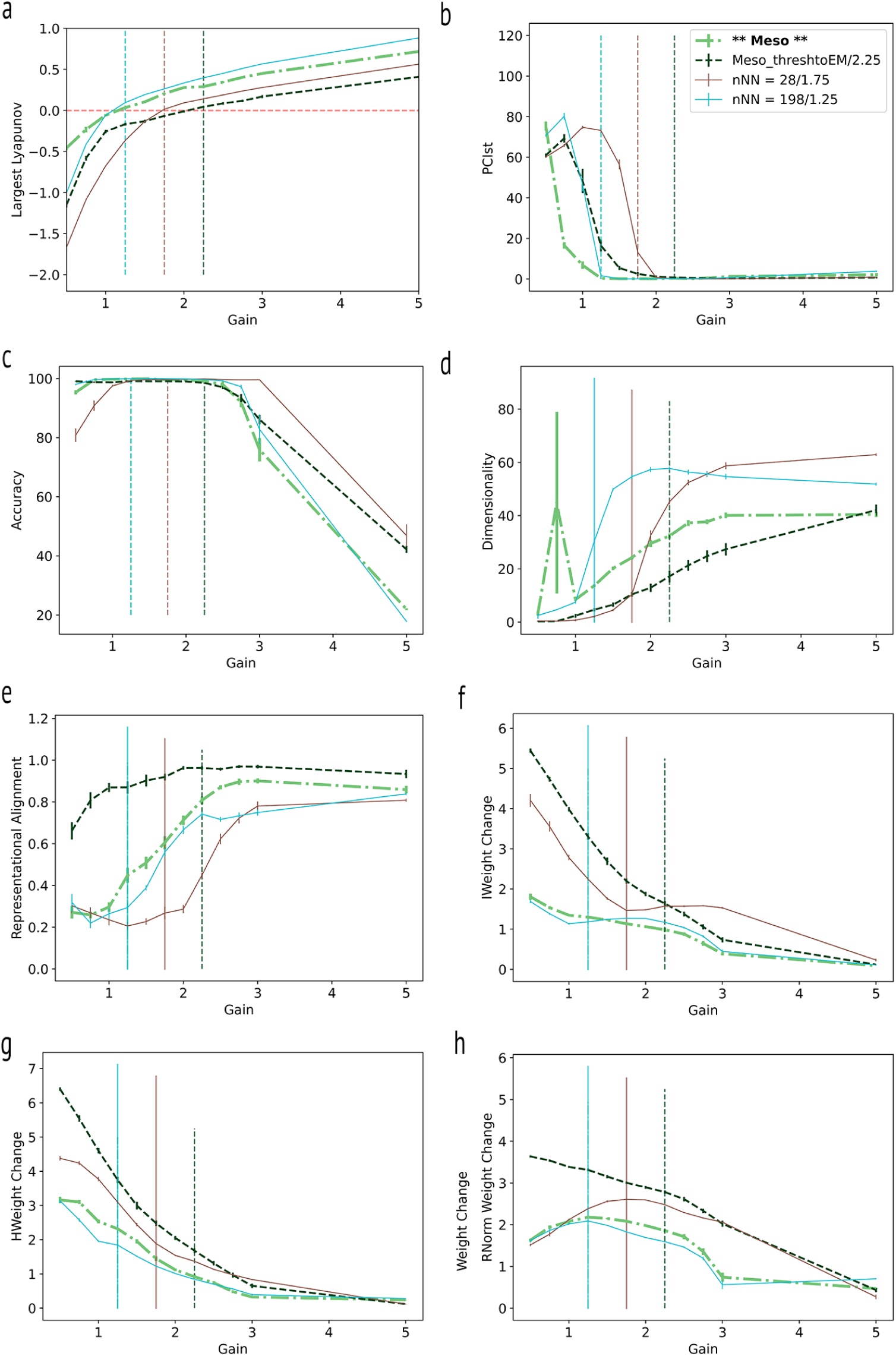
Mesoscopic Connectivity. **(a)** Largest lyapunov exponent *λ* of pre-trained models. True mesoscopic connectivity (indicated as **Meso** in green dash-dot) has a shallower slope than the Gaussian nNN = 198 model(brown). The same holds true for the thresholded mesoscopic model (dark blue - dashed) with respect to the sparsity-matched Gaussian nNN = 28 model (brown) **(b)** Initial PCIst in pre-trained models. **(c)** Model accuracy after 100 epochs of training **(d)** Kaplan-York (KY) dimensionality of trained models **(e)** Representational Alignment. **(f-h)** Norm Weight change in input, hidden and readout layers.

For models based on the cortical column connectivity, the effect of initial gain on dynamical regime is even more pronounced (Figure 7b (V1 23 4 Dales - purple dash-dot), where the maximum Lyapunov exponent is less than 1 for all gains except gains of 1 and 1.25. Moreover, as the gain increases, we observe the maximum Lyapunov exponent decreases rather than increases, deviating substantially from the Gaussian models as well as that of the mesoscopic model.

**Figure 7:**
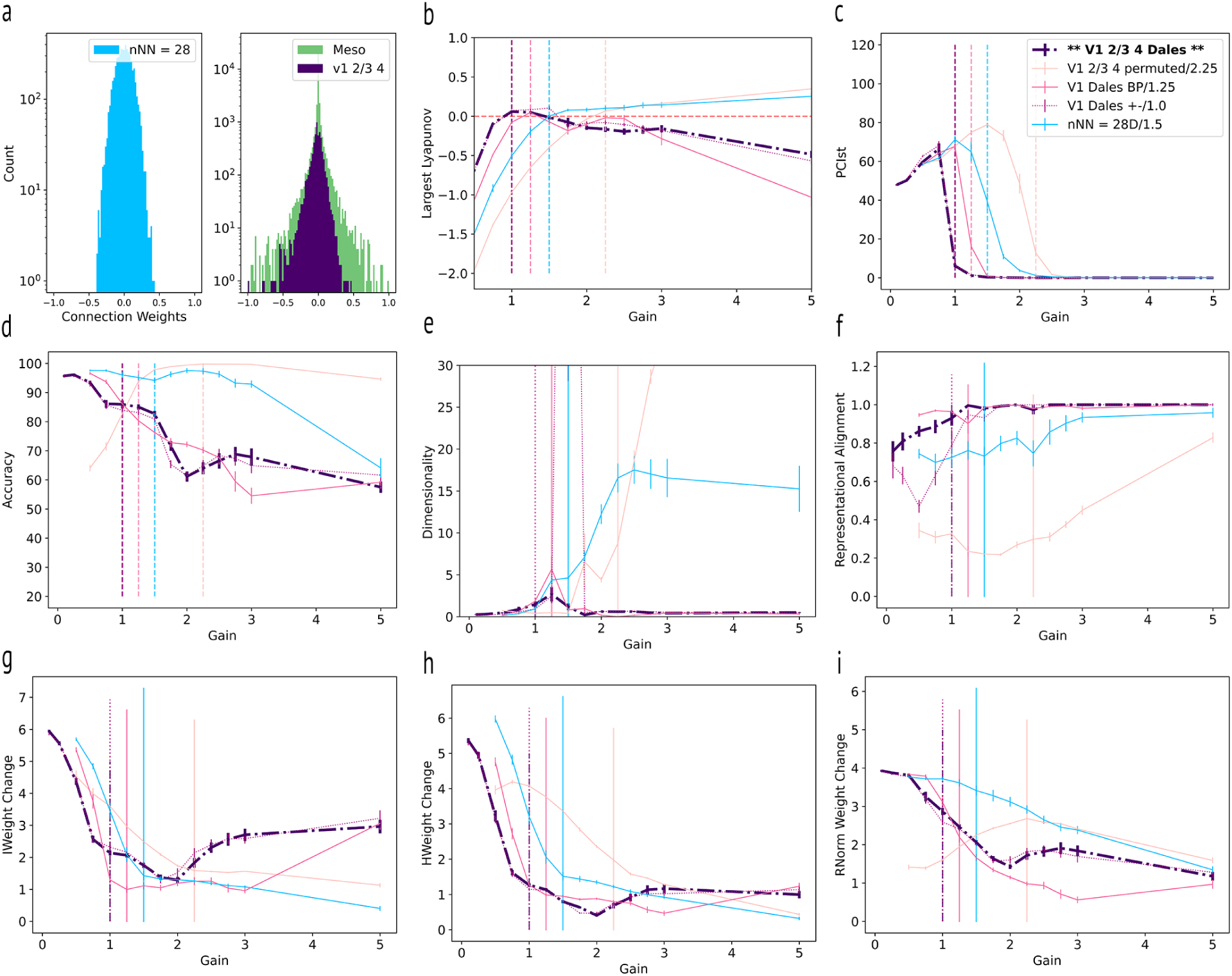
Cortical Column Connectivity. **(a)** Initial distributions of hidden layer connectivity with g = 1.0 for models with nNN = 28 drawn from 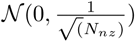 (blue) in comparison to two experimentally derived biological connectivity structures: mouse mesocopic connectiome and synaptic connections within a cortical column of mouse V1. **(b)** Largest lyapunov exponent *λ* of pre-trained models: cortical column connectivity (indicated as ** V1 2/3 4 Dales ** in purple dash-dot), a permuted version of the cortical column in which the distribution is the same but the connection patterns are randomly altered, a block-permuted model of the cortical column where connections are permuted but Dale’s law is preserved, and a sparsity-matched Gaussian nNN = 28 model with Dale’s Law imposed. Biologically realistic connectivity distributions biases these models away from highly chaotic dynamics and lazy learning regime even at high gains. **(c)** PCIst in pre-trained models. **(d)** Model accuracy after 100 epochs of training **(e)** Kaplan-York (KY) dimensionality of trained models **(f)** Representational Alignment. **(g-i)** Norm Weight change in input, hidden and readout layers.

We endeavored to explore which of the characteristics of the cortical connectivity structure (weight distribution, degree distribution, topological structure, Dale’s law, relative balance of excitation and inhibition) enables such stability. To this end we created a series of models with altered structures and found that the observed stability required the combination of block structured connectivity, multiple cell types and adherence to Dale’s law. Below we describe each of the altered structures tested.

1. to test whether Dale’s law alone was sufficient, we created a network in which Dale’s Law was artificially applied to a connectivity structure with Gaussian weights of equal sparsity (Figure 7b - nNN=28D: cyan solid). This model did not exhibit stability at high gains, indicating that Dale’s law alone was not sufficient.
2. to test whether the distribution of weights alone was sufficient, we permuted the cortical column weights such that all topological structure including Dale’s law is disrupted (Figure 7b - V1 2/3 4 permuted: peach). This model also did not exhibit stability at high gains, indicating that weight distribution alone was insufficient.
3. to test whether the degree distribution alone could account for the observed stability, we created a model with Gaussian weights but matched the degree distribution (DD) of the cortical column (Figure S5b - DD green). Degree-distribution alone was not sufficient to maintain stability.
4. to test whether the specific topology of connections was sufficient to reproduce the stability observed in the cortical column, we created a model with Gaussian weights, maintaining the topology (location of non-zero weights) of the cortical column model. Topology alone was also insufficient to maintain stability (Figure S5b - (V1 T magenta solid).
5. To test whether stability could be explained by the combination of weight distribution and topology, we created a network in which the weights are fully permuted but the topological structure is maintained, such that the location of non-zero weights are preserved but Dales law was not preserved (Figure S5b - V1 2/3 4 permuted T, pink dash-dot). This model did result in lower slope as a function of gain, suggesting that the combination of topological structure and weight distribution contributes to stability. However this model did not reproduce the shape of the curve we observed in the cortical column model, where the maximum Lyapunov exponent decreased with increasing gain.
6. To test the possibility that having multiple populations of neurons (E - excitatory, I- inhibitory) with different distributions could account for our observations, we created a model in which cortical column weights were permuted within blocks (E-E, I-I, E-I, I-E), such that block structure and Dale’s law were preserved, but the topological structure was not (Figure 7b - V1 Dales BP: magenta solid). This model did reproduce the stability of the cortical column and the shape of the curve as a function of gain, suggesting that block structure contributes importantly to the stability of the cortical column.
7. a model with weights permuted within block while maintaining topological structure and Dales law (Figure S5b - V1 dales BPT: magenta dash-dot) also maintained stability.
8. Finally, to test whether the overall balance of excitation and inhibition was critical to stability, we created a model in which the signs of all connections are flipped, such that the topology, block structure and Dale’s law are all maintained but the balance of excitation and inhibition is reversed (Figure 7c V1 Dales +-: fuchsia-dotted). Surprisingly, we found that this model was equally as stable as the true connectivity at high gains.

From these models we can conjecture that none of the weight distribution, topology, degree distribution nor balance of excitation and inhibition *alone* is sufficient to reproduce the stability of the cortical column model. Rather, models that matched the observed pattern of stability in the cortical column model featured the combination of a block structured connectivity matrix with multiple cell types and adherence to Dale’s law. We further explored this possibility through a series of simulations (See Supplementary Section 5.4.1), which further suggested that the dynamical stability we observed can be achieved whenever the connectivity contains at least 2 populations of cells, with differing means, such that for each, the population mean is large enough compared to the variance. The simulations also reveal that strict adherence to Dale’s law is not required; approximate adherence to Dale’s law is sufficient. In this case, the mean activity drives the overall system dynamics. Stability is achieved by interaction between excitatory and inhibitory populations in combination with a saturating non-linearity, such that strong oscillations dominate, effectively quenching the chaotic dynamics.

#### 1.5.2 Model Performance

The mesoscopic connectivity models performed similarly to the Gaussian nNN = 198 models to which they are most similar, in that models achieved similar accuracy as a function of gain after 100 epochs (Figure 6c). This is perhaps unsurprising given the matched degree of sparsity and balance of positive and negative weights in both sets of models. Models initialized with connectivity of the cortical column, however, did not perform as well as their nearest Gaussian comparator in terms of sparsity (nNN = 28) (Figure 7d - V1 2/3 4 purple dash-dot trace). Instead, these models only achieved equivalent accuracy to their Gaussian counterparts at much lower gains (*g <* 0.5). Note that Dale’s Law was enforced during training such that the sign of each connection was not allowed to change with back-propagation. Both outcomes are consistent with previous work on networks with cell-type specific connectivity. Specifically, it is known that networks that obey Dale’s law perform more poorly [76], presumably because the constraint that each row of the hidden weight matrix must be either positive or negative, adversely restricts the space of possible solutions. Furthermore, at least one study found that the effective gain of a network with block connectivity structure is greater than the average gain, leading to larger learning capacity at lower gains [77].

#### 1.5.3 PCIst

PCIst, when computed on biologically realistic connectivity models, was similar to that of the previously described Gaussian models. That is, the value increased as a function of initial weight gain, is maximal at the edge of chaos and decreases sharply as the maximum Lyapunov exponent nears zero. However, unlike the Gaussian models (Fig 2d), as the gain is further increased, the maximum Lyapunov exponent becomes negative again for the cortical column model (Figure S13)a. In this case, the PCIst metric reflects the fact that there are nevertheless outliers beyond the unit circle in the eigenspectrum of the connectivity matrix at higher gains (See Supplementary Section 5.4.1). The oscillatory dynamics that dominate at larger gains are amplified enough in the baseline condition as to make the relative response to perturbation negligible.

#### 1.5.4 Learning Regime

In comparison to the Gaussian models, those with mesoscopic connectivity have similar respresentational alignment curves, transitioning towards lazier learning in the chaotic regime. Accordingly, the pattern of weight changes as a function of gain were also similar to the Gaussian models (Figure 6e-h). One exception is the thresholded mesoscopic model, which has larger changes in the hidden weight layer than the other models and higher representational alignment. The difference for this model is likely due to having thresholded the smallest weights while leaving the tails of the distribution, with larger weights. The cortical column models were unique in that they consistently found low-dimensional solutions, despite having notably higher representational alignment for all gains. The apparently lazier strategy employed to reach the solutions in this case likely reflects the restricted solution space of biologically unrealistic training with back-propagation under the constraint of Dale’s law (Figure 7f-i).

## 2 Methods

### 2.1 Model Construction

We trained a series of recurrent neural networks (RNNs) to perform the ten digit (0-9) sequential MNIST task, in which networks sequentially receive 28 rows of 28 pixels from a handwritten digit. The model’s task is to learn to correctly identify the digit after receiving the last row of pixels. All models consisted of an input layer with 28 units (*U*), a single recurrent layer with 198 hidden units (*H* = [198, 198] matrix), and a linear readout layer with input size 198 and output size (*o_t_*) of 10.

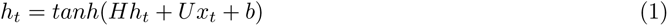

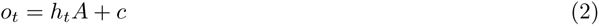

The size of the hidden layer was chosen for consistency with the size of the experimentally derived mesoscopic connectivity of the mouse brain, which reflects the biological connectivity density between 198 major brain areas [74, 75]. The hidden layers of each network were initialized according to a Watts-Strogatz [65] connectivity rule over a range of nearest neighbors (nNN: 4, 8, 16, 28, 32, 64, 128, 198 = fully connected), and rewiring probabilities (p: 0.0, 0.2, 0.5, 0.8, 1.0), while self-connections were prohibited. The resulting networks systematically varied in both their degree of sparsity (nNN = 4 most sparse; nNN = 198 least sparse) and their degree of small-world structure (p = 0.0 highly structured small world architecture; p = 1.0 random Erdös Rényi connectivity). At initialization, non-zero weights of the hidden layer were sampled from a Gaussian distribution with mean *µ* = 0 and standard deviation 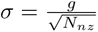 where *N_nz_* is the number of non-zero elements and gain (g : 0.5, 0.75, 1.0, 1.25, 1.5, 1.75, 2.0, 2.25, 2.5, 2.75, 3.0, 5.0), thus normalizing the variance across models at initialization. For each combination (nNN, p, g) we trained 10 randomly initialized models, 4800 models in total.

#### 2.1.1 Biologically Realistic Connectivity

##### Mesoscopic Connectivity

We made use of a previously published whole-brain model of inter-regional mesoscopic connectivity of the mouse brain. The model is based on hundreds of anterograde tracing experiments in C57BL/6J mice across 12 major brain divisions including isocortex, olfactory areas, hippocampus, cortical subplate, striatum, pallidum, thalamus, hypothalamus, midbrain, pons, medulla and cerebellum, allowing for the creation of a whole-brain connectivity model at the scale of 100 *µm* voxels [74, 75]. Hidden layer connectivity between each of 198 regions are their connection densities. Connection density is defined as the sum of the connection strengths from all voxels in a source region to all voxels in the target region divided by the product of the region sizes, where voxel-wise connection strengths represent the fraction of voxel volume expressing fluorescence. Connection density values are positive by definition, but the signs of each connection in the matrix were assigned randomly, making approximately equal number of positive and negative weights. As above, network connection strengths were scaled at initialization by a range of gain values (g : 0.5, 0.75, 1.0, 1.25, 1.5, 1.75, 2.0, 2.25, 2.5, 2.75, 3.0, 5.0). As we are interested in the potential computational advantage of biologically realistic connectivity structure, we did not explore rewiring these model connections (all models p = 0). A sparse mesoscopic connectivity model was subsequently derived from this connectivity matrix by removing connections below a value of .0188 in order to match its sparsity to that of the cortical column model described below.

##### Synaptic Connectivity of a Cortical Column

The hidden layer connectivity of the cortical column model was generated from a data set containing reconstructions of the dendritic trees of hundreds of thousands of neurons as well as their local axonal projections using electron microscopy, giving unprecedentedly accurate information on their 0.5 billion synaptic connections []. The synapses are located within the binocular area of the primary visual cortex (V1) of a single mouse (192 days old). We made use of a subset of this data selecting 198 cells from the set of fully proofread neurons with the nearest euclidean distance from the center point between layers 2/3 and 4. For each neuron in the column, the connectivity strength is calculated as the sum over the volume of each postsynaptic density to each target cell, while noting cell types (excitatory vs inhibitory). For example, if cell a has 10 synapses on to cell b the connection strength of connection *H_ba_* is the sum of the volume of those 10 synaptic densities at cell b. Connections from inhibitory neurons have sign -1, and those from excitatory have sign +1, adhering to Dale’s law [78, 79]. The resulting connection matrix included 52 inhibitory cells and 146 excitatory cells. However, as the inhibitory cells tend to make a larger number of local connections. the ratio of inhibitory to excitatory weights was 1.60. The cortical column models differ substantially in both the distribution of their weights (Figure 7a) as well as their degree of sparsity. The sparsity of the cortical column-derived hidden layer is 85.71%, while the mescoscopic connectivity structure is fully connected, with the exception of self-connections. It is for this reason we included the nNN = 28 model as it matches the sparsity of the cortical column model (Table 1). Once again, networks connection strengths were scaled at initialization by a range of gain values (g : 0.5, 0.75, 1.0, 1.25 1.5, 1.75, 2.0, 2.25, 2.5, 2.75, 3.0, 5.0) and for these models rewiring probability p = 0.

### 2.2 Training

All networks were trained using the Adam optimizer for 100 epochs with a learning rate of 1e-5, a batch size of 256 and cross-entropy loss. Note that while connection variance across models was normalized at initialization, it was not controlled over training. Model readout occurred at a single time point immediately following input of the last row of sMNIST digit input. Hidden states were initialized as zero. Sparsity was enforced over learning such that only non-zero weights were updated over the course of training. For cortical column models with cell-type specific connectivity, Dale’s law was enforced over training, such that sign changes were prevented by clamping negative weights to a maximum of -.0001 and positive weights to a minimum of .0001.

### 2.3 Identifying Chaotic and Ordered Learning Regimes

#### 2.3.1 Lyapunov Exponents

Gradients can explode or vanish exponentially over recurrent steps through a network, especially when the connection strengths between recurrent processing units are large, making models numerically unstable and difficult to train. As Lyapunov exponents represent the exponential growth rates of nearby trajectories in phase space of the model, we compute the finite time Lyapunov spectrum as a measure of model stability. Following the standard QR-decomposition technique [80] for computing the Lyapunov spectrum, we compute the average, over input samples, of the eigenvalues of the Jacobian of hidden state dynamics of each model using the python implementation in [66].

Specifically: A matrix Q is initialized as the identity matrix and hidden states are initialized as zero. At each of the T = 28 time points through the model, we compute the Jacobian matrix (first-order partial derivatives of equation 1 with respect to the hidden state dynamics h) and the product of the Jacobian matrix with the Q matrix. QR decomposition is applied to this product and used to update Q, which tracks the relative expansion or contraction of the model over time. The Lyapunov exponent of the *i^th^* batch at timestep t *r^i^* is the expansion of the *i^th^* vector corresponding to the *i^th^* diagonal element of R, and the *i^th^* Lyapunov exponent *λ_i_* is then given by

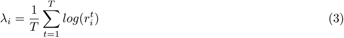

The exponents *λ_i_* are computed in parallel for a batch of input samples (size 100) of sMNIST test data to drive the networks and are subsequently averaged over the batch. We compute Lyapunov exponents both at network initialization and after training. In Figure 2a,b, we report the maximum Lyapunov exponent resulting from this process after averaging over each of 10 model instantiations of a given model parameter combination (nNN, p, g) for p = 1.0. (See Figure S1 for similar results obtained for rewiring probabilities [0, 0.2, 0.5, 0.75]). Chaotic dynamics are indicated by positive average largest Lyapunov exponent (*λ >* 0). Though we computed the spectrum both pre and post-training, we define the transition from ordered to chaotic *g_cnNN_* dynamics as the gain at which the largest Lyapunov exponent *λ >* 0 in the network prior to training.

### 2.4 Assessing Sensitivity to Noise

Model sensitivity to noise was assessed by adding Gaussian noise (0, 0.01) to sMNIST test data. We compare the accuracies of the trained model (trained in the absence of noise) to test data with and without noise. Models with chaotic dynamics are expected to exhibit greater sensitivity to noise and therefore greater decreases in accuracy.

### 2.5 PCIst

PCIst is a measure of the spatio-temporal complexity of the evoked response of a system to perturbation [35, 38]. It was developed for use with EEG data in response to a perturbation delivered via transcranial magnetic stimulation (TMS). However, the metric is applicable more generally to any evoked signal composed of a baseline state and a well-defined response period to a perturbation in a network of causally interacting units; thus, importantly it can be applied to artificial networks. Computing PCIst involves several steps: 1) Baseline and response periods are defined. 2) Singular value decomposition (SVD) is performed on the matrix consisting of the response time-series of the hidden states over time 3) Principal components are selected from the eigenvalues of the decomposed matrix so as to account for a user-specified amount of variance. 4) components are then selected in terms of their signal-to-noise ratio (SNR), calculated as the square root of the ratio of average response power 5) recurrence quantification proceeds on the remaining components by computing a distance matrix between all time points 6) This matrix is thresholded at several values and transition matrices are computed as the number of times the state crosses each threshold. The ”st” in PCIst is derived from quantifying the number of state transitions in this segment of the analysis, where state transitions are measured. 6) An optimal threshold *ɛ* is determined such that the number of state transitions in the response relative to baseline is maximized. 7) The average number of state transitions in the baseline and response period of the n^th^ component is the difference in the number of state transitions in the *ɛ*-thresholded matrix of the response relative to the baseline scaled by the number of response samples. 8) PCIst is then the sum of these differences over components. Therefore, PCIst is simply a product of the number of retained components (reflecting the spatial differentiation of the response across the network), and the average number of state transitions across components (which reflect the temporal complexity present in each component.) See [38] for more details.

In our study, the time series used to compute PCIst comes from the hidden state space of the model. We defined the response period as the 28 steps corresponding to the trajectory of the network to a single sMNIST test digit (the perturbation). Since the network’s response unfolds over 28 time steps, we defined a baseline period of equal length during which the network receives a small Gaussian noise input sampled from (0*, .*01). The perturbation was a batch (size 256) from the sMNIST test data set of a given digit. The metric was computed on the mean over batches of the hidden layer states for baseline period and perturbations. PCIst was computed on all networks both at initialization (IPCIst) and after training on the sMNIST task.

### 2.6 Dimensionality

Dimensionality was computed from the Lyapunov spectrum of the trained models, as the attractor dimension using the Kaplan-Yorke conjecture [81]. First the Lyapunov spectrum was computed as in 2.3.1 on the trained rather than pre-trained models. The resulting average values for a particular model parameter set (nNN, p, g) are sorted from largest to smallest. Then let j be the largest index for which the sum of the cumulative sum of the exponents is greater than zero.

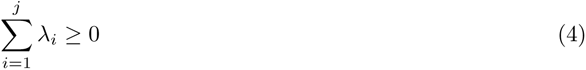

and

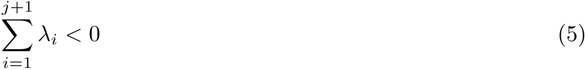

Then the dimension

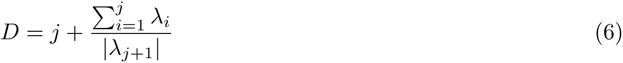

### 2.7 Characterizing Learning Regime

#### 2.7.1 Weight Change

To assess the magnitude of weight changes at each layer of the network, we computed the *L*^2^ or euclidean norm of the difference between the final weights *W ^f^* and the initial weights *W* ^0^.

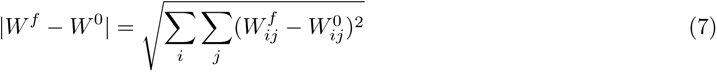

#### 2.7.2 Representation Alignment

Representation alignment (RA) quantifies the directional change in the representational similarity matrix (RSM) due to training. Instead of focusing on the dissimilarity as used in representational similarity analysis [82], RSM focuses on the similarity between how two pairs of input are represented by computing the Gram matrix of last step hidden activity.

An increased level of representation alignment indicates a higher degree of lazy learning in the network, and it is obtained by [16]:

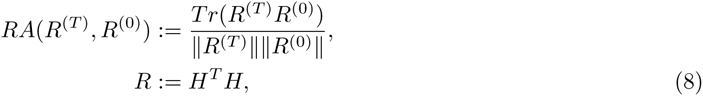

where *H* is the hidden activity, and *R*^(0)^ and *R*^(*T*)^ are the initial and final RSM, respectively.

#### 2.7.3 Tangent kernel alignment

Tangent kernel alignment provides a measure of the directional change in the neural tangent kernel (NTK). The NTK is a mathematical tool that calculates the inner product of gradients for each input pair. Like representational alignment (RA), tangent kernel alignment calculates the Gram matrix between input pairs. However, unlike RA, it does so based on the gradient rather than the final hidden activity, thereby quantifying network similarity in terms of the gradients. In more specific terms, the NTK, for any given pair of inputs, determines the covariance between the gradients of the neural network’s output with respect to its parameters.

A heightened degree of tangent kernel alignment points to a greater extent of lazy learning within the network. As outlined in [16], this is calculated as follows:

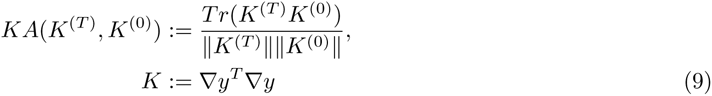

where *K*^(0)^ and *K*^(*T*)^ denote the initial and final NTK, respectively.

## 3 Discussion

In this study, we centered our attention on recurrent neural networks (RNNs) and revealed that the manipulation of initial hidden weight gain results in ”lazier” or ”richer” learning. In the latter, which is brought about by smaller initialization gains, we observe more significant network changes (including substantial weight alteration and rotation of representation) as well as lower-dimensional representation. These findings are in line with existing literature on feedforward networks [14, 69]. To the best of our knowledge, prior research specifically characterizing the dichotomy of rich and lazy learning within the context of RNNs is scant [83]. Expanding on this exploration, we ventured further into the interplay between the rich and lazy learning transition and its connection to the transition to chaos. Our investigation characterizes solution attributes and learning behavior on both sides of this transition and suggests that rich representations can be biased into a network’s learning strategy by tuning the Lyapunov spectrum towards ordered dynamics.

In the context of infinite-depth feedforward networks, the correlation between the rich/lazy transition and the chaotic/ordered transition has been previously documented [84, 85]. However, this exploration has yet to be extended to RNNs. Given the divergent behavior at the infinite-depth limit, and if such an infinite-depth limit in feedforward networks translates to an infinite sequence length limit in RNNs, our study admittedly has some limitations. Notably, we only considered finite sequence lengths in this work, leaving the exploration of infinite sequence limits as an area for future inquiry. Additionally, more nuanced categorization of learning regimes is left for future work [83]. Further, we manipulated the initial weight scale to tune between the learning regimes, but we recognize there are other parameters — such as network width, scaling of readout weights (*α* parameter) [14, 16] and initial weight rank [71] — that could also be adjusted to affect the transition. Hence, future research will explore these additional dimensions.

Additionally, in this work, we explored differences in network dynamics and learning regime on biologically realistic connectivity structures at two different scales. These structures differ in their weight distributions, degree distributions, inclusion of Dale’s law, and the balance of positive/negative weights. In the mesoscale model, we found that longer-tailed distributions with only a single population did afford greater stability over a wider range of gains than Gaussian models, while learning equally well. Importantly, using the cortical column model, we found that having multiple populations from different distributions changed network dynamics dramatically. Theoretical work has yet to fully describe the expected dynamical transitions for connectivity matrices with multiple populations drawn from separate distributions, each with different non-zero means and standard deviations. Our results are nevertheless consistent with previous theoretical work showing that block connectivity structure with more than one population will have outliers in their eigenspectrum, when the sum of synaptic weights into each neuron is non-zero, while noting that such networks can have non-intuitive dynamics [86].

Finally, we computed PCIst, a clinically relevant measure of consciousness, for these models. To our knowledge, this is the first time that the metric has been characterized in artificial neural networks in relation to their associated Lyapunov exponents. Interestingly, we found that the metric can be predicted by the value of the maximum Lyapunov exponent, peaking at the edge of chaos, with larger values in the non-chaotic regime where we also observed a tendency towards rich learning. It is important to note that the biological connectivity structures used in our study result from fine-tuning both prenatally and over the course of the first months of life. As a result, their ordered dynamics may reflect the impact of prior tuning through biological feedback mechanisms to have this desirable characteristic. The result suggests that 1. biological systems are biased towards rich learning. 2. consciousness as measured by PCIst may be an evolutionary consequence of favoring rich learning strategies.

Although beyond the scope of this work, our observation that networks shift towards lazier learning strategies as dynamical complexity increases, raises interesting questions about brain states associated with more complex neural activity, such as when under the influence of psychedelics. The use of psychedelics in the treatment of psychiatric disorders has become increasingly commonplace. Although their therapeutic mechanism is not understood, recent research has found that psychedelics robustly increase brain complexity [87–89]. It has been suggested that such treatment may increase brain flexibility [90, 91]. Thus, this work raises an intriguing, testable hypothesis that presumed increased flexibility may include a shift in learning strategy under the influence of these drugs.

Overall, our work connects critical network dynamics, learning regimes, and measures of consciousness and characterizes the influence of network sparsity, structure, and weight variance on network dynamics and learning strategy. We show that both learning regime and measures of consciousness undergo a transition at the coupling strength that delineates chaotic from ordered dynamics. As we continue to investigate, these findings promise to unlock deeper understanding and more robust applications within the field of artificial intelligence and neuroscience.

## Acknowledgements

We wish to thank the Tiny Blue Dot Foundation, the NSF and the NIH for their support as this work was funded in part by grants from the Tiny Blue Dot Foundation, NSF 2223725, and NIH R01EB029813 and RF1DA055669. We thank Guillaume Lajoie and Stefano Recanatesi for their valuable discussion and feedback on this work. We also wish to thank the Allen Institute founder, P. G. Allen, for his vision, encouragement and support.

## 4 Supplementary

### 4.1 Maximum Lyapunov Exponents

We reported the maximum Lyapunov Exponents for models with rewiring probability = 1.0 (Erdös-Rényi) in the main text. Here we report similar results for all other rewiring probabilities [0.0, 0.2, 0.5, 0.8]. The overall pattern of transitions into the chaotic regime prior to training, with largest Lyapunov exponent *λ >* 0 are consistent with the Erdös-Rényi connectivity models, such that the sparsest models transition at higher gains and the least sparse models transition at lower gains (Figure S1 - left column). Additionally, for all models, *λ* tunes closer to the critical point (*λ* = 0) with training, regardless of whether *λ* is positive or negative prior to training (Figure S1 - right column).

### 4.2 Dimensionality sMNIST 2-digits

We trained an additional series of models for rewiring probability = 1.0 only, on the 2-digit sMNIST task, using digits [2,5] and found that models exceeded KY dimension of 2 rather than 10 at *g_cnNN_* (Figure S3).

### 4.3 Neural Tangent Kernel

The NTK provides another method for quantifying the richness/laziness of learning [15]. The NTK increases towards a maximum value of one as network models begin to use lazier learning strategies. Consistent with the other metrics used to quantify the richness/laziness of learning, for most models, the NTK increases as a function of gain as networks approach the transition to chaos (Figure S4). However, because it is based on the gradients of the network’s output with respect to its parameters, values can become numerically unstable as the model dynamics become chaotic. We nevertheless, include them for completeness.

### 4.4 Biologically Realistic Connectivity

We endeavored to identify the characteristics of the cortical connectivity structure (weight distribution, degree distribution, topological structure, Dale’s law, relative balance of excitation and inhibition) that led to negative maximum lyapunov exponents at high gain. As described in the main text, we tested many altered connectivity structures to see which aspect of connectivity was necessary to achieve this. Here we show several additional models described but not shown in the main text (Figure S5.)

#### 4.4.1 Generative Model Reproducing Dynamical Stability of Cortical Column Model

Biological neural networks can be divided into various sub-populations of morphologically and functionally defined cell types. Here, we focus on the two most basic cell types: excitatory and inhibitory. We split neurons in our experimental data set into two sub-populations ((E)xcitatory and (I)nhibitory). Experimentally estimated synaptic weights can be rewritten as a block matrix of the form

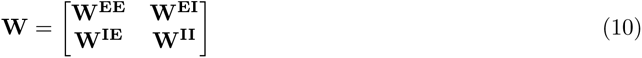

In order to generate random surrogate weights we compute statistics (means and variances) of weights between each possible pair of populations (*E E*, *E I*, *I E*, *I I*). More precisely, for a given pair of populations (*α, β*) of sizes (*N ^α^, N ^β^*), we take the set of non-zero synaptic weights from neurons in population *β* to neurons in population *α* and calculate the mean 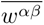 and variance 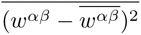. We define *µ^αβ^* as the sum of mean weights that, on average, any neuron from population *α* receives from neurons in population *β*. If there are *C^αβ^* non-zero *β → α* weights, the average number of non-zero weights from population *β* per neuron in population *α* is *K^αβ^* = *C^αβ^/N ^α^* and we should have

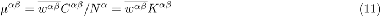

Similarly, we define *α^αβ^* as

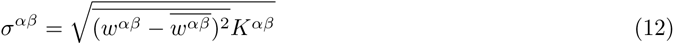

**Table S1:** Model Critical Points.

| nNN | p | Transition point |
| --- | --- | --- |
| 4 | 0.00 | 5.0 |
| 4 | 0.20 | 3.0 |
| 4 | 0.50 | 2.5 |
| 4 | 0.75 | 2.5 |
| 4 | 1.00 | 2.5 |
| 8 | 0.00 | 2.75 |
| 8 | 0.20 | 2.25 |
| 8 | 0.50 | 2.25 |
| 8 | 0.75 | 2.25 |
| 8 | 1.00 | 2.25 |
| 16 | 0.00 | 2.25 |
| 16 | 0.20 | 2.0 |
| 16 | 0.50 | 2.0 |
| 16 | 0.75 | 2.0 |
| 16 | 1.00 | 2.0 |
| 28 | 0.00 | 1.75 |
| 28 | 0.20 | 1.75 |
| 28 | 0.50 | 1.75 |
| 28 | 0.75 | 1.75 |
| 28 | 1.00 | 1.75 |
| 32 | 0.00 | 1.75 |
| 32 | 0.20 | 1.75 |
| 32 | 0.50 | 1.75 |
| 32 | 0.75 | 1.75 |
| 32 | 1.00 | 1.75 |
| 64 | 0.00 | 1.5 |
| 64 | 0.20 | 1.5 |
| 64 | 0.50 | 1.5 |
| 64 | 0.75 | 1.5 |
| 64 | 1.00 | 1.5 |
| 128 | 0.00 | 1.25 |
| 128 | 0.20 | 1.25 |
| 128 | 0.50 | 1.25 |
| 128 | 0.75 | 1.25 |
| 128 | 1.00 | 1.25 |
| 198 | 0.00 | 1.25 |
| 198 | 0.20 | 1.25 |
| 198 | 0.50 | 1.25 |
| 198 | 0.75 | 1.25 |
| 198 | 1.00 | 1.25 |

The surrogate network is fully connected and is not subject to strict Dale’s law. Its weights form a block matrix with entries generated randomly from normal distribution with matched experimental statistics. The numbers of neurons in each population, *N*°^E^ and *N*°*^I^*, do not have to match those in experimental data. Means and variances are rescaled by the number of neurons in the presynaptic population. More precisely, the random surrogate weight matrix takes the form

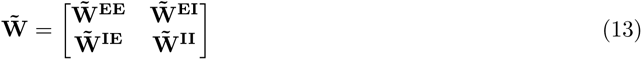

**Table S2:** Connectivity statistics of the subset of EM V1 Cortical Column data set used in our numerical experiments.

| $\alpha\beta$ | $K^{\alpha\beta}$ | $\overline{w^{\alpha\beta}}$ | $\mu^{\alpha\beta}$ | $\sigma^{\alpha\beta}$ |
| --- | --- | --- | --- | --- |
| EE | 4.8<br>(0.25) | 0.061<br>(0.002) | 0.290<br>(0.018) | 0.117<br>(0.005) |
| EI | 17.0<br>(0.56) | -0.058<br>(0.0012) | -0.992<br>(0.038) | 0.248<br>(0.009) |
| IE | 27.9<br>(3.09) | 0.054<br>(0.0013) | 1.51<br>(0.17) | 0.261<br>(0.019) |
| II | 18.2<br>(1.05) | -0.083<br>(0.0036) | -1.52<br>(0.011) | 0.467<br>(0.033) |

where entries of each block are generated i.i.d. as

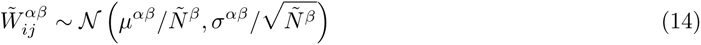

We simulated networks driven entirely by recurrent connections (i.e., without any inputs) with either experimental or surrogate weights (Figure S8). Although the eigenvalues of the resulting random matrix do not match eigenvalues of experimental weights (Figure S8b,f), the qualitative features of its dynamics, including suppression of chaos (Figure S8a,e) and low-dimensional, periodic or quasiperiodic attractors (Figure S8c,d,g,h) are reproduced. In the surrogate network, the emergence of oscillations is driven by the presence of a pair of extreme eigenvalues (”outliers”). The position of the outliers can be predicted (Figure S8f) from the eigenvalues of the 2 *×* 2 matrix:

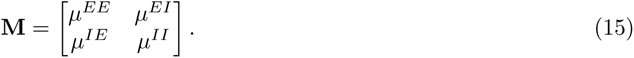

The effect of chaos suppression disappeared when standard deviations of weight distributions were increased by a factor of 2 (Figure S8i-l). These results were not finite-size effects as confirmed in simulations of larger surrogate networks (Figure S9).

Overall, our findings suggest that the main drivers of the chaos-suppressing oscillations in the original network may be imbalanced excitatory and inhibitory input weights in tandem with relatively low variability of the weights around the mean. However, the experimental eigenspectrum is markedly more complex than the eigenspectrum of random surrogate weights (Figure S8b,f), indicating that our simple generative model does not capture other, potentially crucial features of the original weight matrix. Outliers could for example appear due to pairwise weight co<u>rrela</u>tions or cell-type-specific connectivity

The estimated mean weights 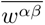 had comparable magnitudes for all four types of connections and the differences in the values of *µ^αβ^*, although statistically significant, were driven mostly by large differences in the values of *K^αβ^* (see Table S2). In particular, *K^EE^* was much smaller than *K^EI^*, leading to *µ^EE^ < µ^EI^*. At this point it is worth noting that the EM V1 Cortical Column data is focused on local circuits as it does not include projections to a given neuron from neurons from outside its near neighborhood. This raises the possibility that the lack of E/I balance in the original network may only reflect local circuit connectivity patterns. In this view, statistics at larger (”global”) spatial scales may be significantly different and we may expect the overall connectivity to be closer to the balanced regime. Importantly, however, the presence of multiple complex-valued outliers in the mean-balanced regime is still possible, albeit only if mean weights strongly dominate over their standard deviations (*µ^αβ^ ≫ σ^αβ^*) [92], see Figure S10. Indeed, due to the local circuit sampling, the experimental values of *K^αβ^* in the data set are much lower than the total average number of presynaptic partners per neuron. Since the ratio *µ^αβ^/σ^αβ^* scales like 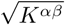 (assuming that global and local connectivity statistics are comparable), this may be the regime the underlying biological networks are operating in.

#### 4.4.2 NTK on Biologically Realistic Connectivity

We report on the NTK results of the mesoscopic and cortical column models in the main text. Additionally, for completeness, we have included the results for all other model variants tested (Figure S11).

#### 4.4.3 Relationship between PCIst and Lyapunov Exponents

In all models we explored, we found a similar relationship between the maximum Lyapunov exponent and PCIst. However, the size of the models we explored were kept constant. We therefore explored two larger models with recurrent layers of size 500 and 1000. From these larger models we see that the point at which PCIst begins to decrease moves closer to the edge of chaos as the size increases (Figure S12).

As we observed for Gaussian networks, for biologically realistic connectivity, PCIst drops off sharply when the Lyapunov exponent becomes positive (Figure S13). As gain increases further, the maximum Lyapunov exponent becomes negative again without a corresponding increase in PCIst. In this regime, where (as described above) chaos is quenched by oscillatory dynamics, the metric reflects the outliers in the eigen-spectrum beyond the unit circle and the fact that perturbations to such networks do not result in strong signal beyond the background response to a Gaussian noise input (See Methods).

---

**Figure S1:**
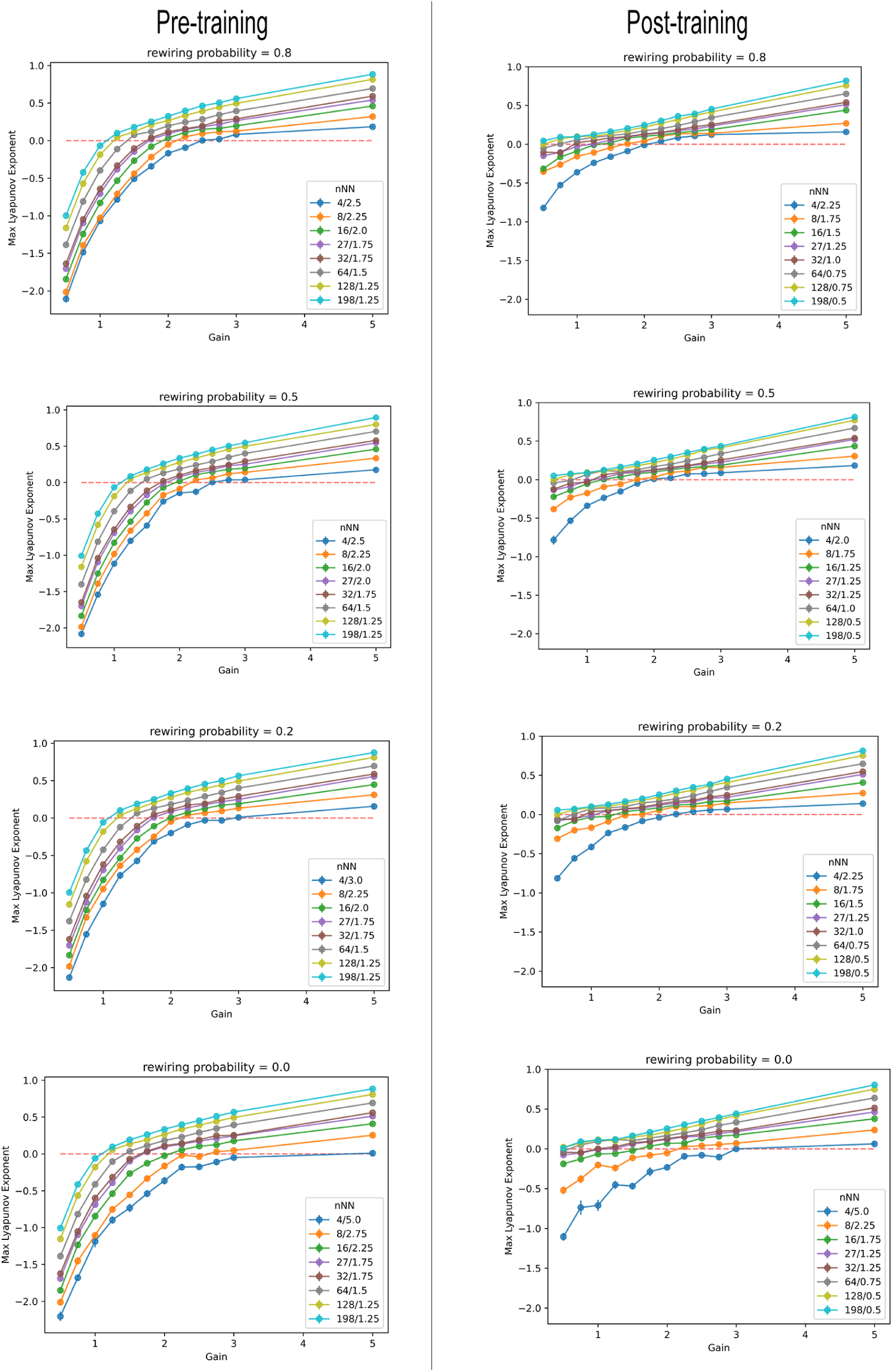
Maximum Lyapunov Exponents for rewiring probabilities [0.0, 0.2, 0.5, 0.8]. Pre-trained models (left column) Post-training (right column).

**Figure S2:**
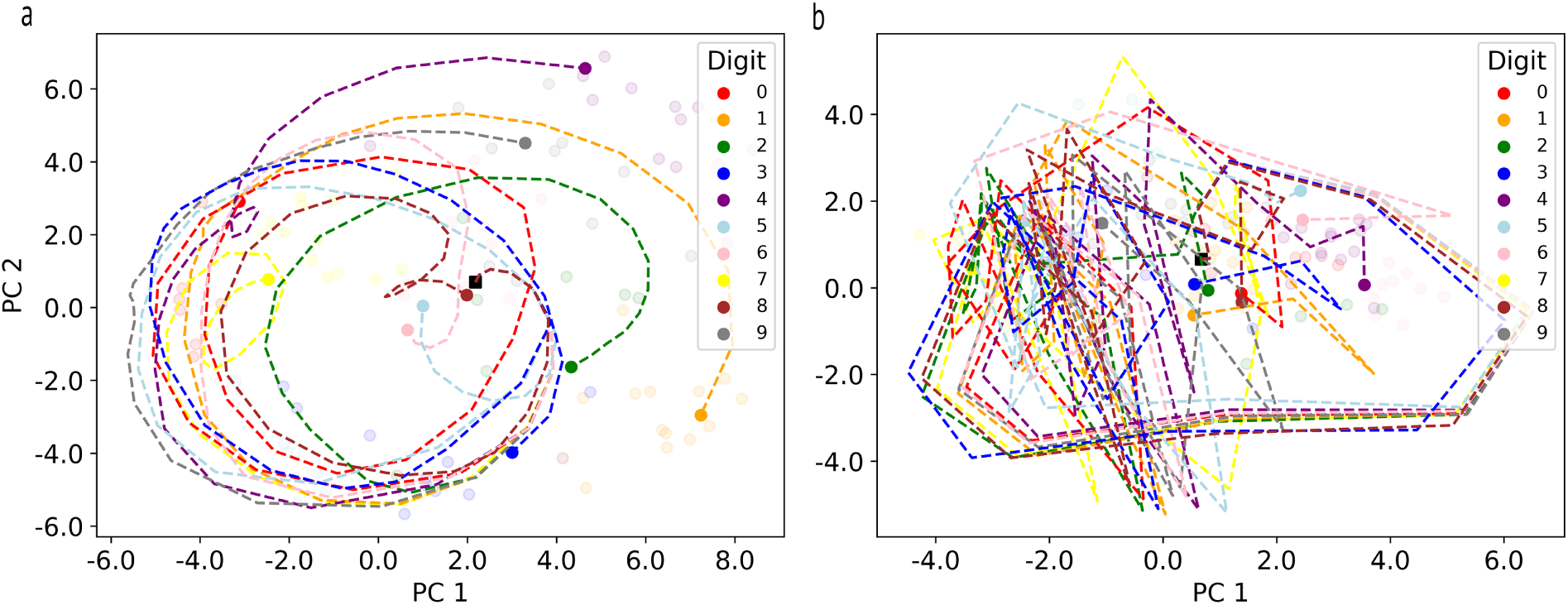
Example Trajectories for nNN = 28, p = 1.0 **(a)** Trajectory associated with ordered learning at gain = 1.0 *< g_cnNN_* projected onto the first 2 principle components of the hidden states. Colors indicate MNIST digit class. **(b)** Trajectory associated with chaotic learning at gain = 2.5 *> g_cnNN_*

**Figure S3:**
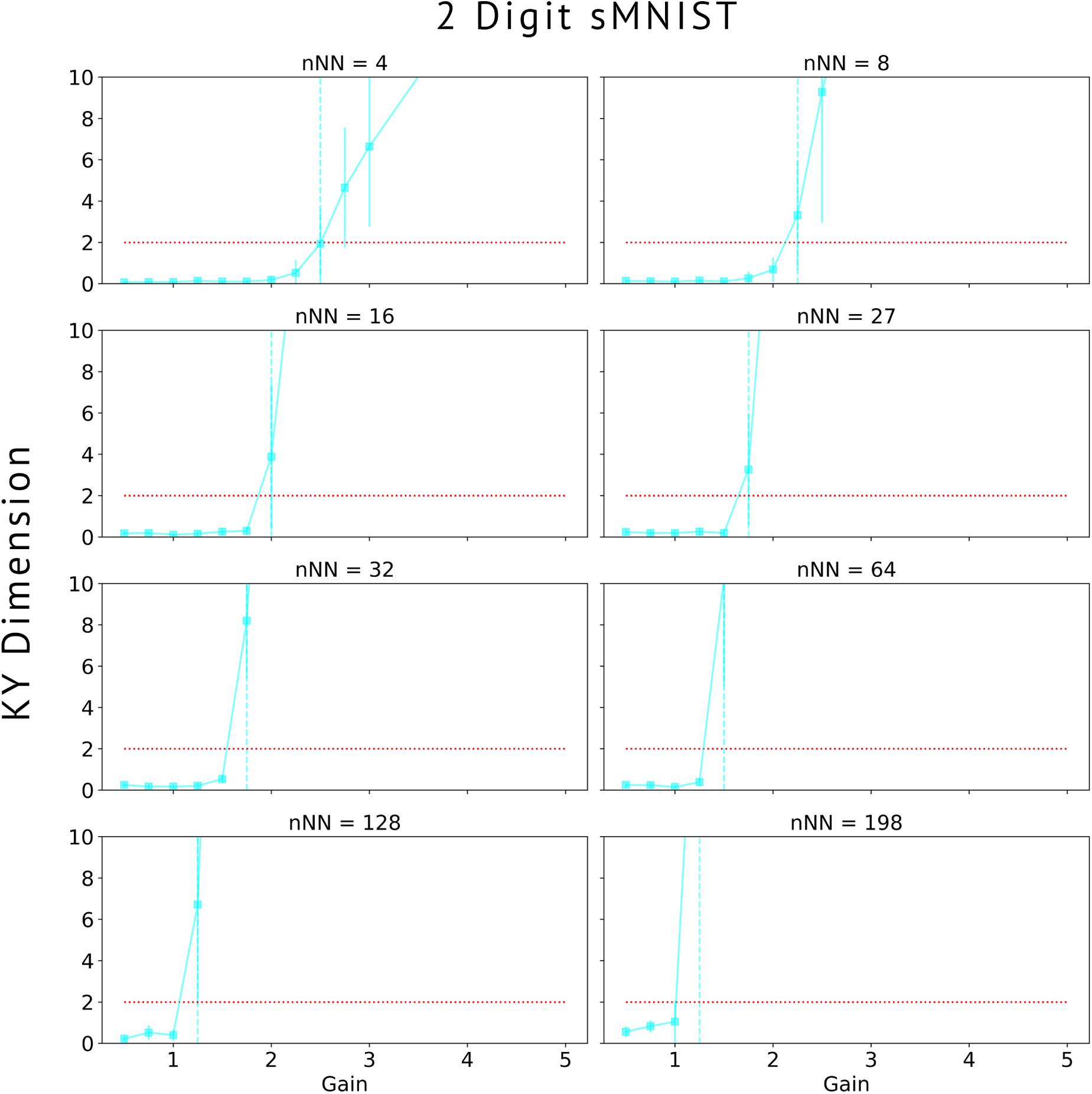
KY dimensionality for the 2-digit sMNIST task. Dotted red line = two.

**Figure S4:**
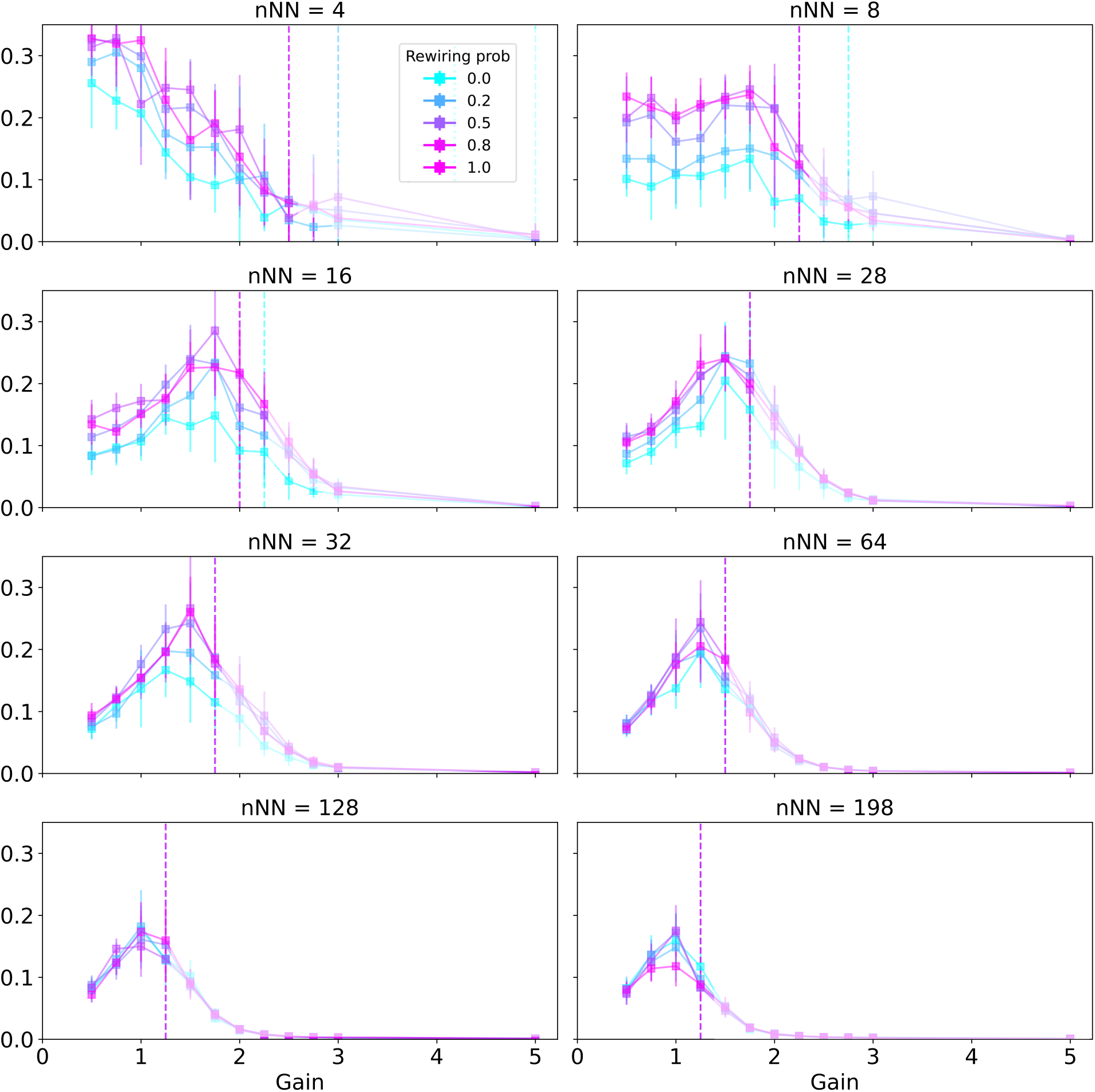
Neural Tangent Kernel Alignment. NTK values after the transition to chaos are greyed to indicate the fact that they may reflect numeric instabilities in their estimation.

**Figure S5:**
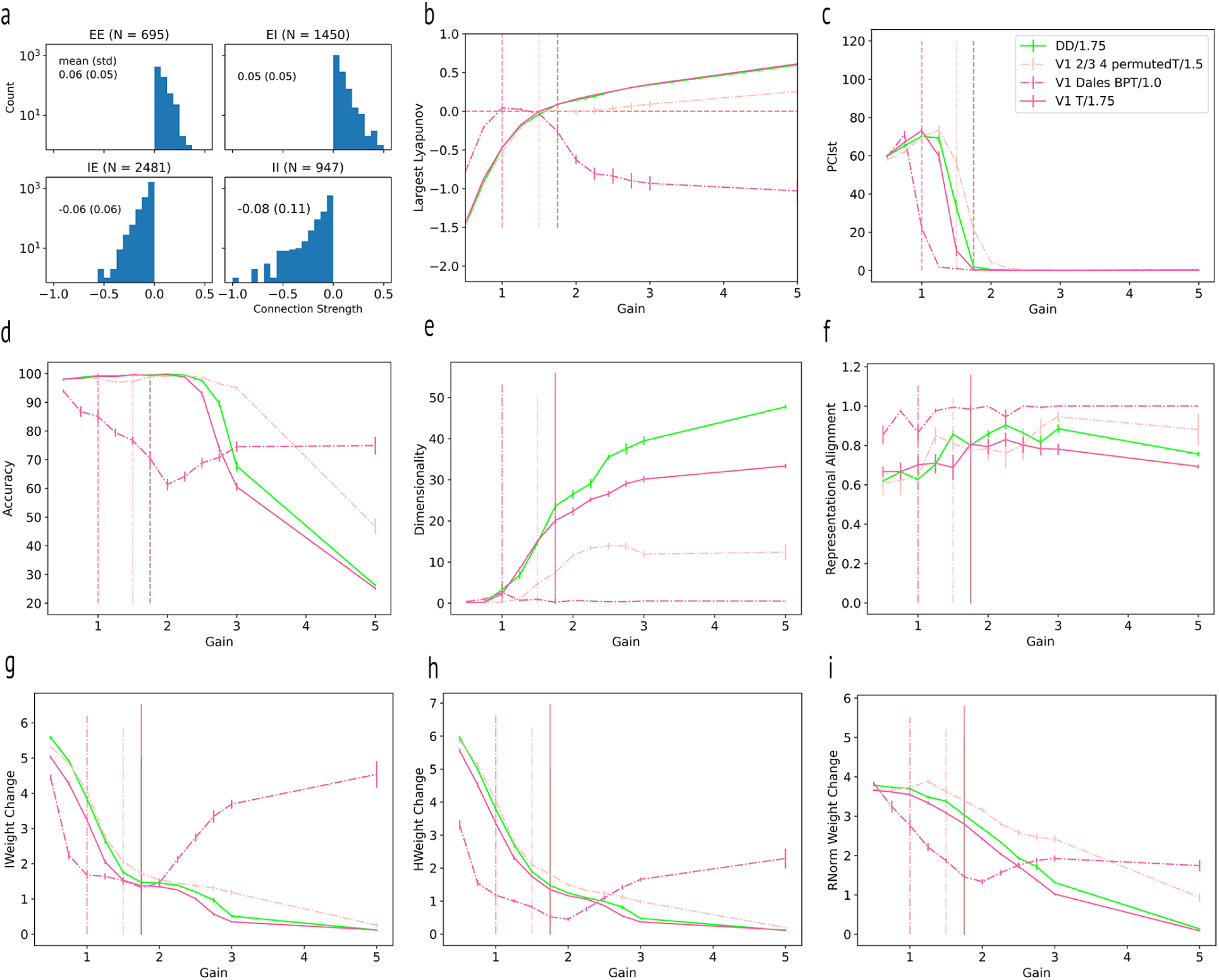
Cortical Column model and variants **(a)** Cortical column - distribution of weights for each cell type (E/I) block (E-E), (E-I), (I-E), (I-I) connectivity. **(b)** Maximum Lyapunov Exponents **(c)** PCIst pre-training **(d)** Accuracy achieved after 100 epochs of training on sMNIST task **(e)** KY Dimensionality of the final state. **(f)** Representational alignment **(g)** Input layer norm weight change **(h)** Hidden layer norm weight change **(i)** Output layer norm weight change

**Figure S6:**
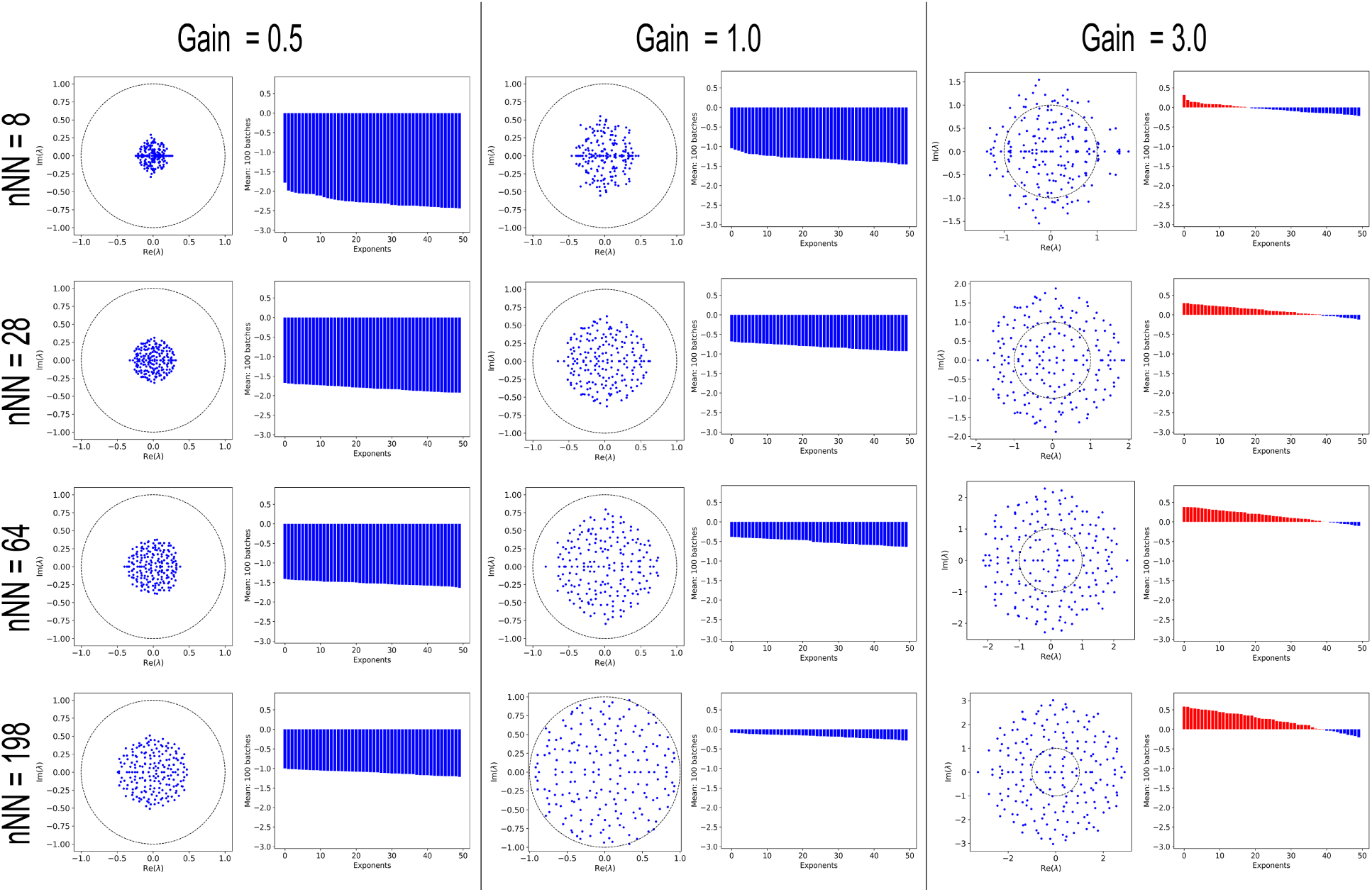
Lyapunov Exponents and Eigenvalues for Gaussian Models

**Figure S7:**
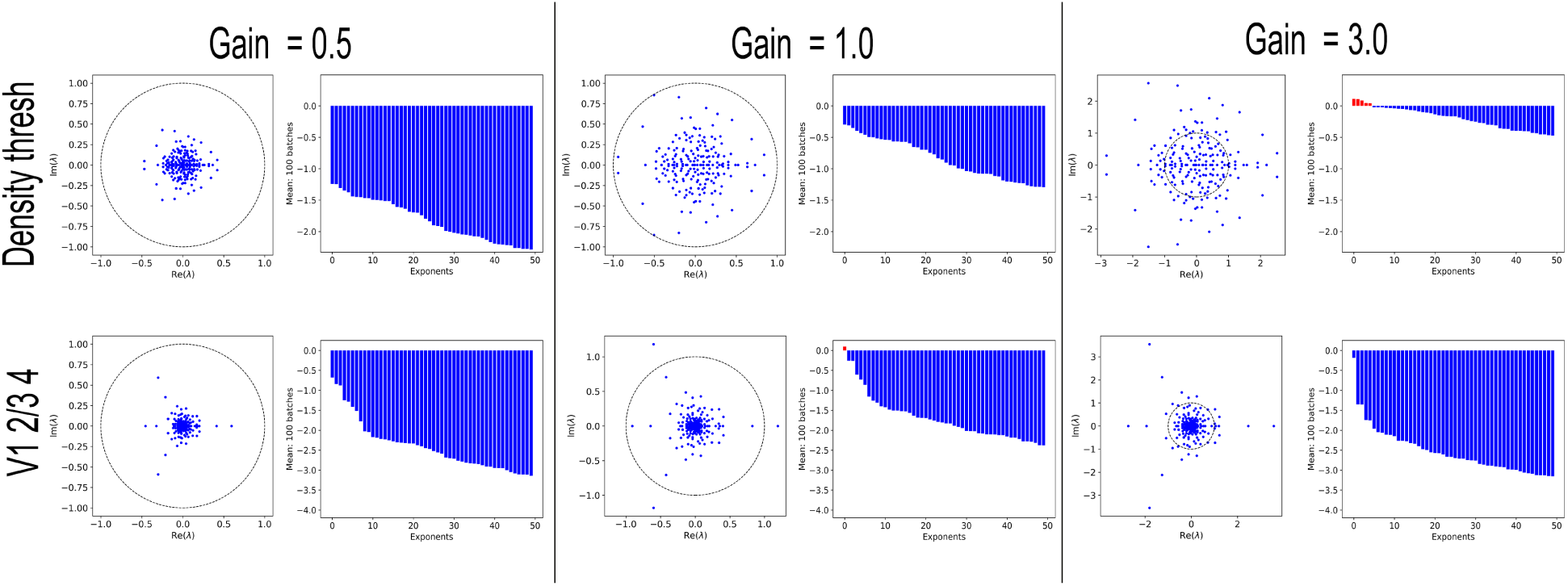
Lyapunov Exponents and EigenValues for Biologically Realistic Connectivity Models

**Figure S8:**
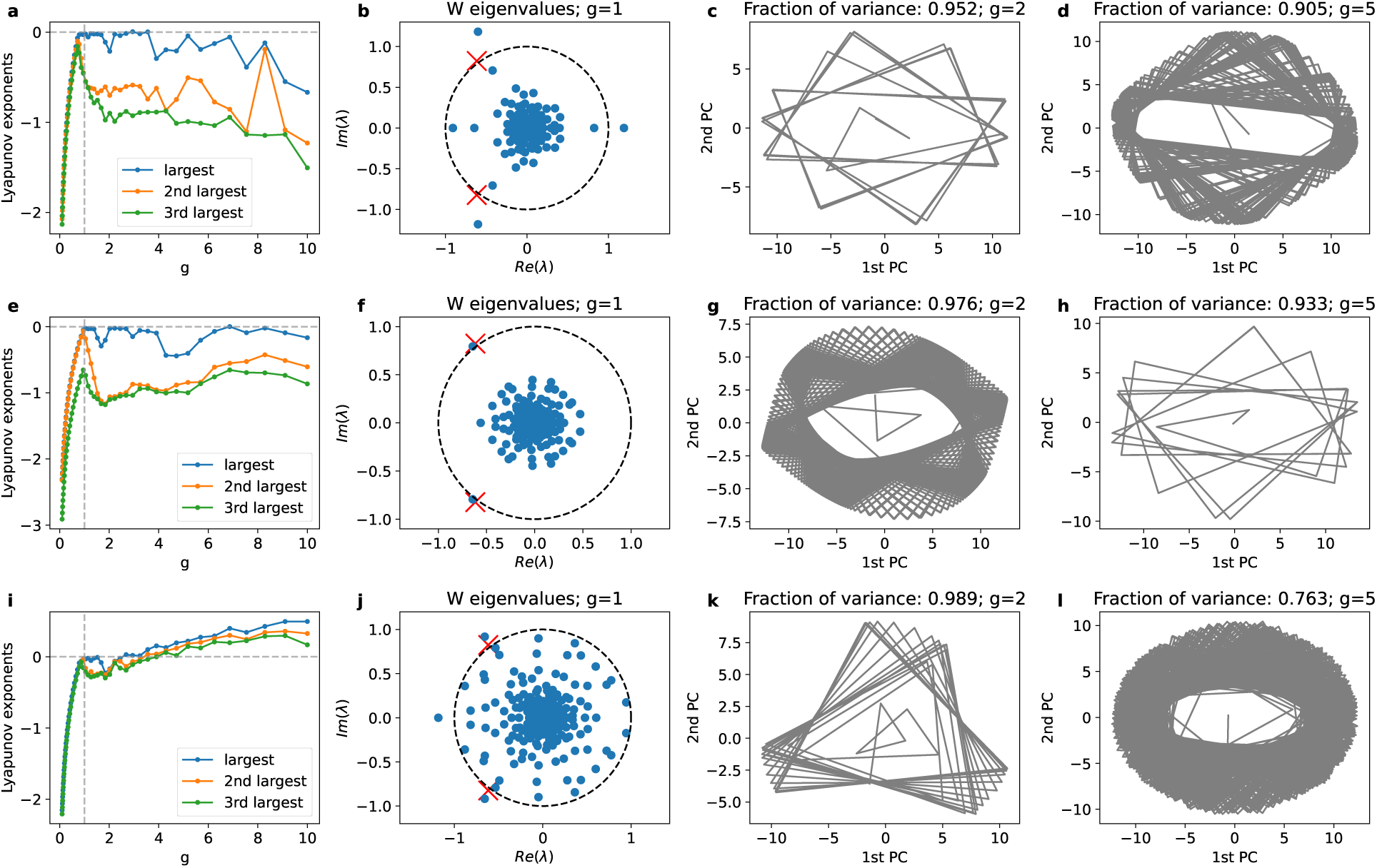
Random surrogate weights can reproduce qualitative features of neural dynamics driven by experimental weights. **(a)** Three largest Lyapunov exponents as functions of gain *g* using experimental weights. **(b)** Eigenvalues of the experimental weight matrix (*g* = 1). Most eigenvalues are contained within a relatively small circular-shaped core, but the presence of multiple outliers suggest non-random features of the connectivity. Red crosses correspond to a pair of eigenvalues of a 2 × 2 matrix **M** constructed from experimental mean weights. They do not match any of the outliers very well, but are close to the most extreme pair of complex-valued outliers. **(c)** Trajectory of the first two principal components for *g* = 2. **(d)** Same as (e) but with *g* = 5. **(e-h)** Same as (a-d) but with random surrogate weights with statistics matched to experimental data. The number of neurons in each population is the same as in the experiments, leading to a relatively large realization dependence (not shown), but the qualitative features of chaos suppression is robust. **(i-l)** Same as (e-h) but with *σ^αβ^* scaled up by a factor of 2. Here weight distributions are wide enough to diminish the influence of average inter-population structure. Thus, the classical scenario of transition of chaos is recovered.

**Figure S9:**
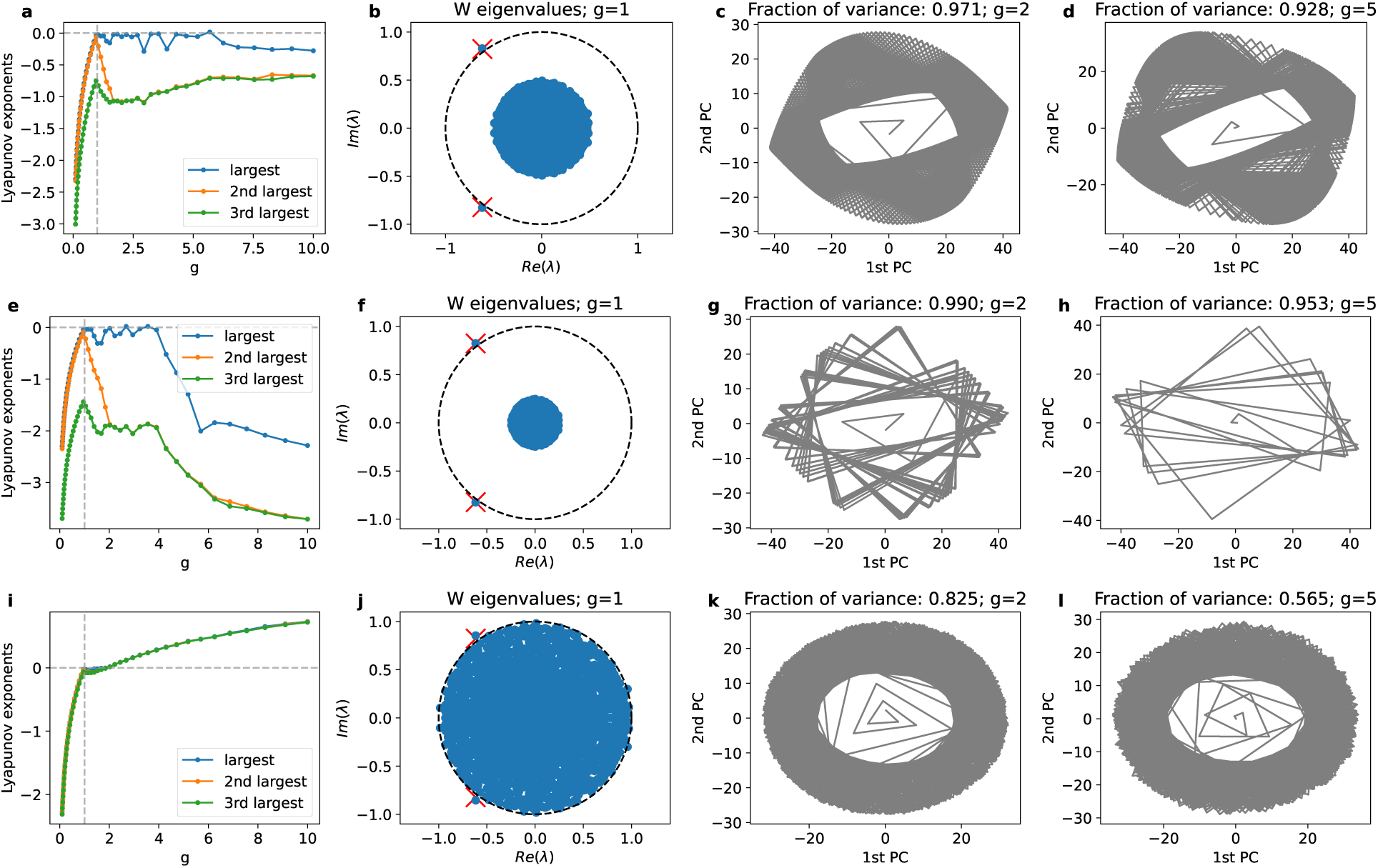
Results obtained with random surrogate weights are qualitatively the same as we increase the number of simulated neurons. **(a-d)** Same as Figure S8(b-c) but with larger populations (*N ^E^* = *N ^I^* = 1000) and appropriately rescaled moments otherwise matched to experimental data. **(e-h)** Same as (a-d) but with *σ^αβ^* scaled down by a factor of 0.5. **(i-l)** Same as (a-d) but with *σ^αβ^* scaled up by a factor of 2.

**Figure S10:**
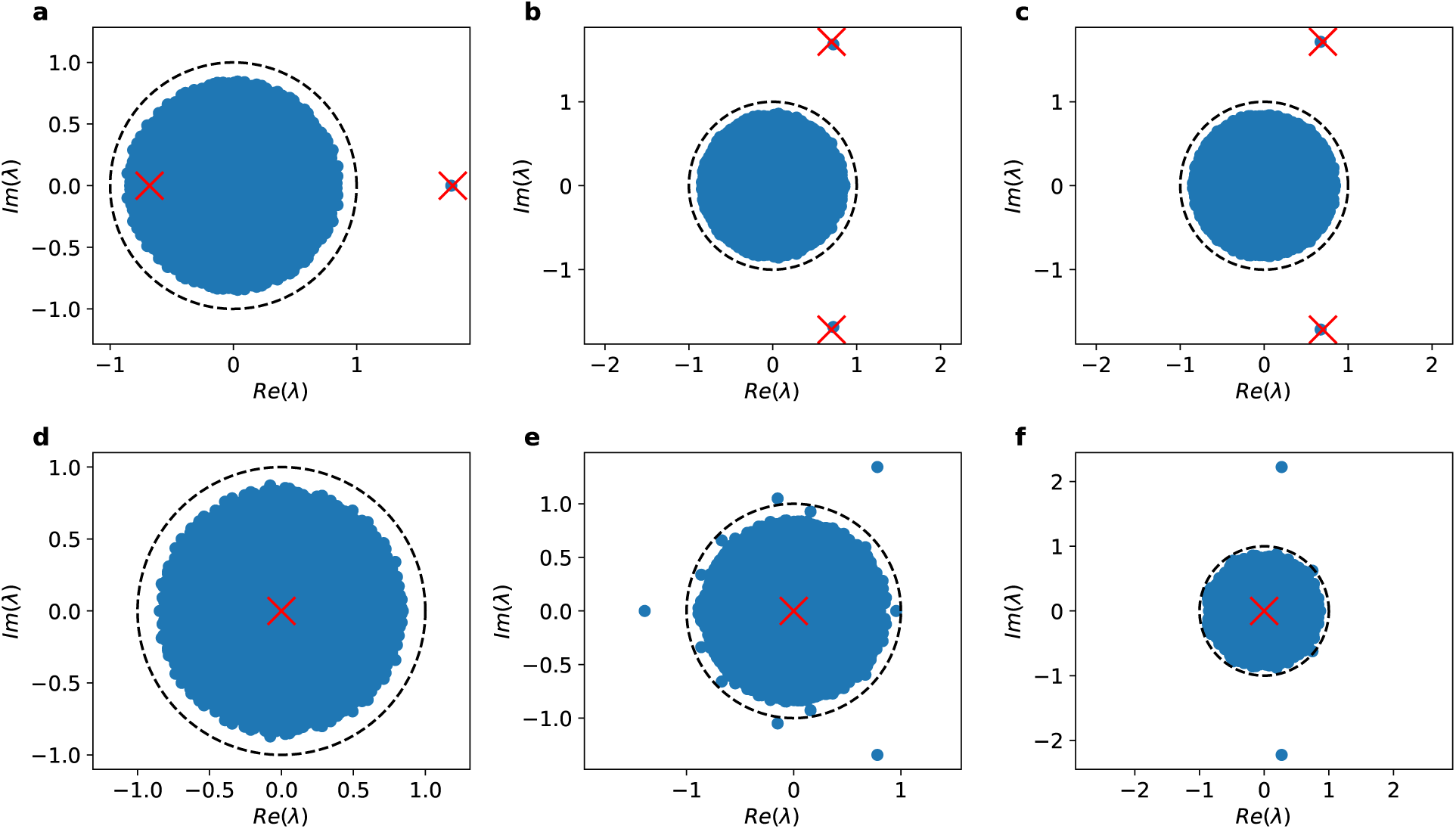
Eigenvalues of the weight matrix **W** depending on the matrix of means **M**. Here, *N ^E^* = *N ^I^* = 1000 and *σ^αβ^* = 0.6. **(a)** 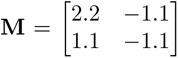. There are two real eigenvalues of **M** but only one lies outside of the disk defined by the circular law, so matrix *W* features a single deterministic outlier. **(b-c)** 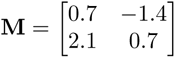. A pair of complex outliers is well-predicted by the eigenvalues of matrix **M**. (b) and (c) correspond to two independent realizations of the weight matrix; the locations of the outliers are subject to small fluctuations that are expected to disappear with *N ^E^* = *N ^I^*. **(d)** 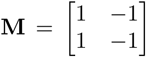. In the balanced regime, **M** has no non-zero eigenvalues and, as a result, no clear outliers are produced. **(e-f)** 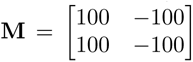. Although **M** has no non-zero eigenvalues, clear outliers are produced as a result of the large magnitude of the low-rank perturbation. The positions of the outliers are not deterministic, as confirmed by comparing two independent realizations in (e) and (f), and as such cannot be directly predicted based solely on **M**.

**Figure S11:**
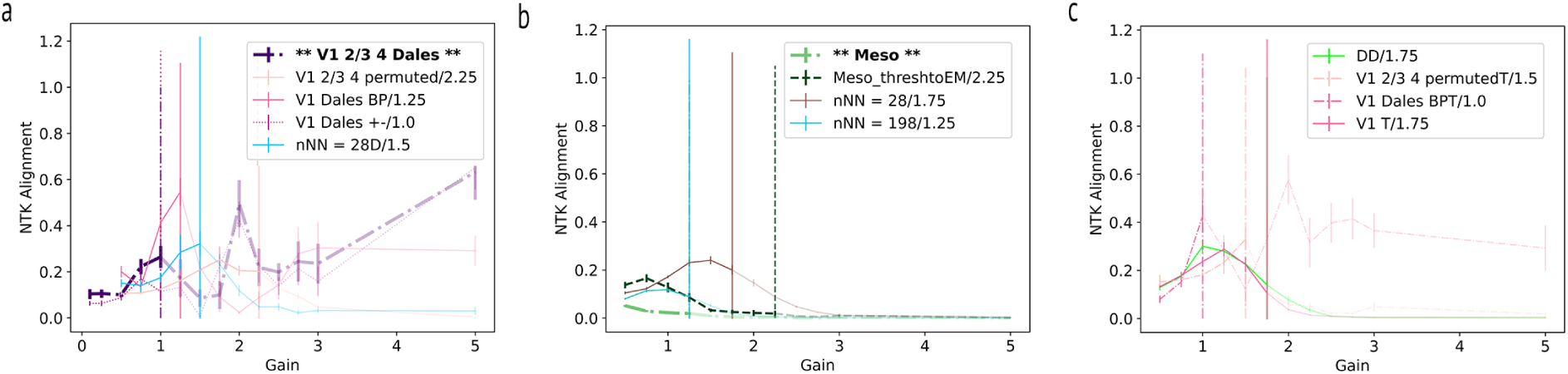
Neural tangent kernel alignment in networks with biologically realistic connectivity.

**Figure S12:**
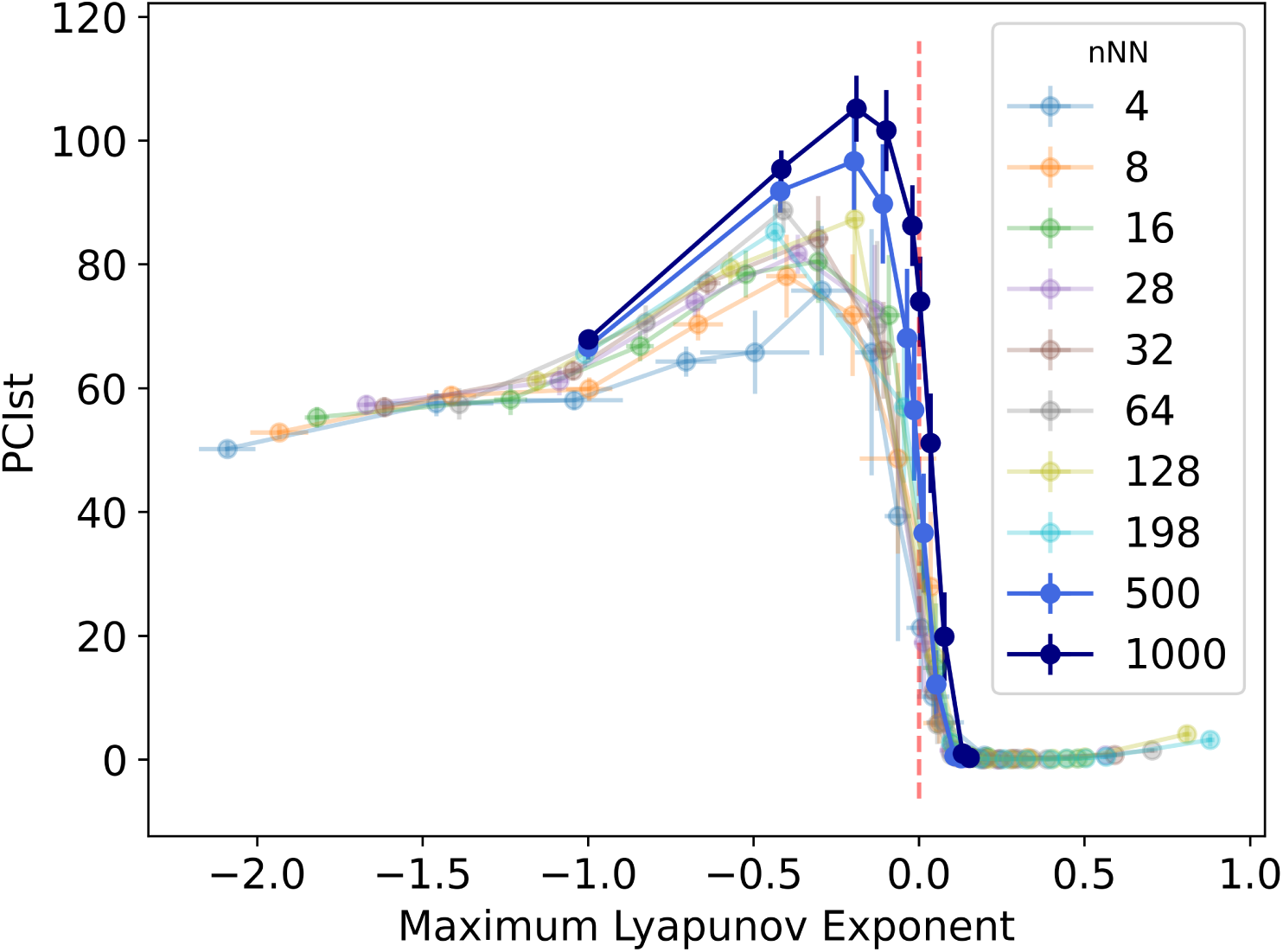
Relationship between PCIst and largest Lyapunov Exponent: Finite Size Effects. The point at which PCIst begins to decrease moves closer to the edge of chaos as networks increase in size.

**Figure S13:**
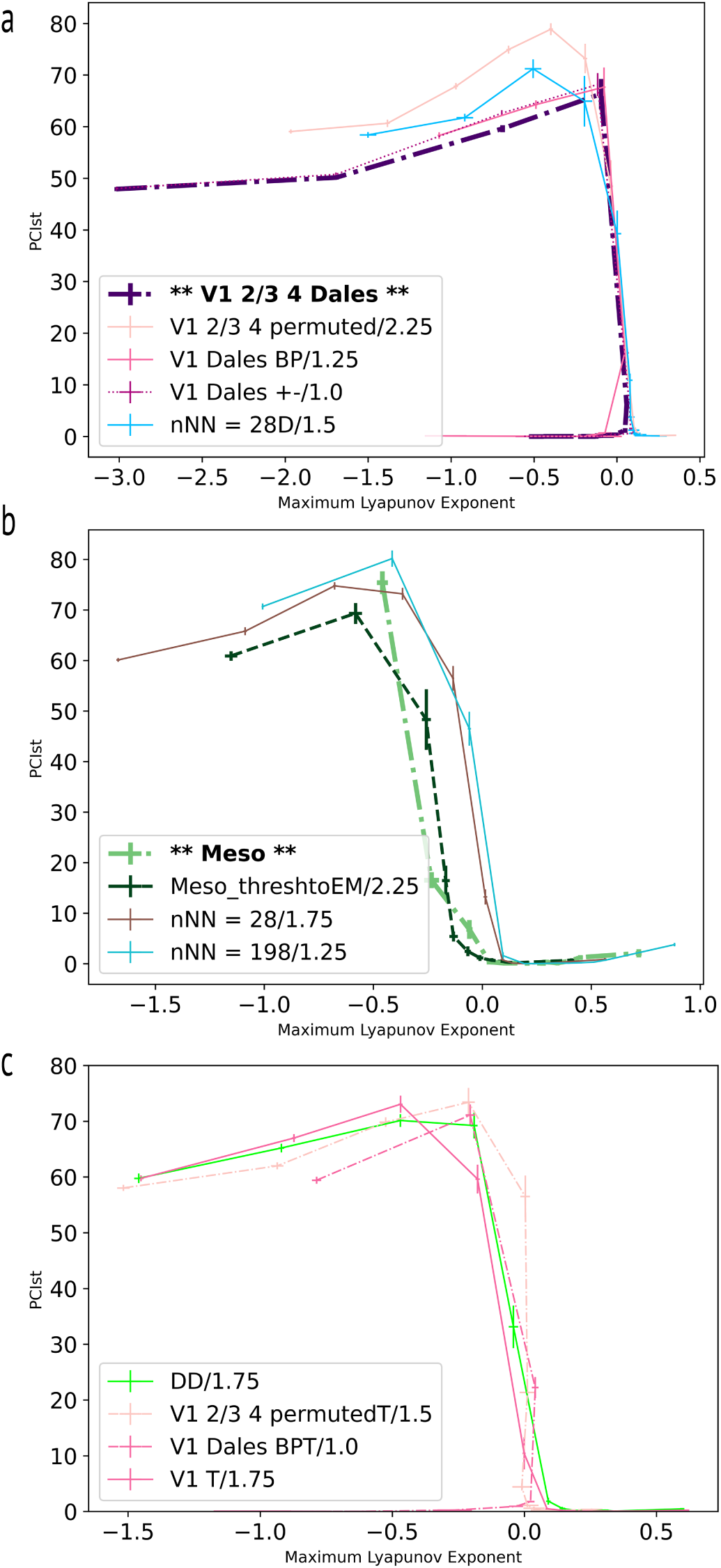
Relationship between PCIst and Lyapunov Exponents in all other model variants tested. Biologically realistic connectivity structure indicated in bold. PCIst decreases towards zero in chaotic regime as well as in models with Dale’s Law with eigenspectrum outliers leading to oscillatory quenched chaos.

